# Absolute Quantification of Viral Proteins from Pseudotyped Vesicular Stomatitis Virus (VSV-GP) using Ultra High-Performance Liquid Chromatography- Multiple Reaction Monitoring (UPLC-MRM)

**DOI:** 10.1101/2023.10.09.561556

**Authors:** Rajeswari Basu, Richard Dambra, Di Jiang, Sophia A. Schätzlein, Shu Njiyang, Joseph Ashour, Abhilash I. Chiramel, Adam Vigil, Vladimir V. Papov

**Author notes:** Corresponding Authors, **Rajeswari Basu**-Materials and Analytical Sciences, Boehringer Ingelheim Pharmaceuticals Inc., Ridgefield, Connecticut 06877;, **Vladimir. V. Papov**, Jr.-, Materials and Analytical Sciences, Boehringer Ingelheim Pharmaceuticals Inc., Ridgefield, Connecticut 06877. **Di Jiang**: Denali Therapeutics Inc., Development Sciences, South San Francisco, CA, 94080.

## Abstract

The rapidly developing field of oncolytic virus (OV) therapy necessitates development of new and improved analytical approaches for characterization of the virus during production and development. Accurate monitoring and absolute quantification of viral proteins is crucial for OV product characterization and can facilitate the understanding of infection, immunogenicity, and development stages of viral replication. Targeted mass spectrometry methods, like multiple reaction monitoring (MRM), offers a robust way to directly detect and quantify specific targeted proteins represented by surrogate peptides. We have leveraged the power of MRM by combining ultra-high performance liquid chromatography (UPLC) with a Sciex 6500 triple stage quadrupole mass spectrometer to develop an assay that accurately and absolutely quantifies the structural proteins of a pseudotyped vesicular stomatitis virus intended for use as a new biotherapeutic (designated hereafter as VSV-GP) to differentiate it from native VSV. The new UPLC-MRM method provides absolute quantification with the use of heavy labeled reference standard surrogate peptides. When added in known exact amounts to standards and samples, the reference standards normalize and account for any small perturbations during sample preparation and/or instrument performance, resulting in accurate and precise quantification. Because of the multiplexed nature of MRM all targeted proteins are quantified at the same time. The optimized assay has been enhanced to quantify the ratios of the processed GP1 and GP2 proteins while simultaneously measuring any remaining or unprocessed form of the envelope protein GPC (full-length GPC).

**IMPORTANCE:** Development of oncolytic viral therapy has gained considerable momentum in the recent years. VSV-GP is a new biotherapeutic emerging in the oncolytic viral therapy platform. Novel analytical assays that can accurately and precisely quantify the viral proteins are a necessity for the successful development of viral vector as a biotherapeutic. We developed a UPLC-MRM based assay to quantify the absolute concentrations of the different structural proteins of VSV-GP. The complete processing of GPC is a pre-requisite for infectivity of the virus. The assay extends the potential for quantifying full-length GPC, which provides an understanding of the processing of GPC (along with the quantification of GP1 and GP2 separately). We used this assay in tracking GPC processing in HEK-293-F production cell lines infected with VSV-GP.

## INTRODUCTION

Knowledge of viral protein expression levels plays an important role in viral vector process development and therefore, a well characterized viral biotherapeutic should include an assay that can monitor the absolute concentrations of the viral proteins in purified drug product (1). The analytical methods employed toward this goal should be precise, quantitative, and reliable to ensure efficient and robust process control during production (2, 3). Proteomics-based methods can prove useful for determining protein expression levels during the virus life cycle (3, 4). While protein expression profiling is now routine in pharmaceutical research, the use of the more targeted MRM approach provides sensitive, accurate, and precise information on viral protein levels (5, 6). MRM is a tandem MS (MS/MS) scan mode that utilizes the unique capability of a triple stage quadrupole (QQQ) mass spectrometer (6–8). Target peptide molecular ions are selected by the first mass analyzer (Q1), fragmented in (Q2) by collision-induced dissociation (CID) with one or several of the resultant fragment ions uniquely generated from the precursor peptide ion then selected by the second mass analyzer (Q3). The precursor-fragment ion pairs are termed MRM transitions that can be measured sequentially and repeatedly at a periodic interval faster than the chromatographic elution of the peptide resulting in chromatographic peaks for each transition which then allows simultaneous quantification of multiple peptides (9–14). Quantification of full-length proteins is accomplished by generating surrogate peptides via enzymatic digestion. Standard solutions are prepared using known amounts of synthetic surrogate peptide and analyzed to generate standard curves for determination of the surrogate peptide concentration in samples. The method can be adapted for absolute quantification by spiking in known amounts of stable isotope (heavy) labeled reference standards with the exact same amino acid sequence as the unlabeled surrogate peptides (15–17). The heavy standard reference spikes normalize for any variability across samples (such as sample injection volume or slight changes in instrument performance) making this method absolute, accurate, and precise (18–20).

The Vesicular Stomatitis Virus (VSV) is used widely both as an oncolytic virus and vaccine vector (21, 22). Herein, we use a variant of VSV that has been pseudotyped with the glycoprotein (GP) from LCMV (23, 24) and apply UPLC-MRM to quantify the levels of all structural proteins encoded by the VSV-GP genome. The 11 kb VSV genome encodes for 5 proteins: nucleocapsid (N), phosphoprotein (P), large polymerase (L), matrix (M) and surface glycoprotein (GP) serving different functionalities that have been well studied (24–29). Protein composition was compared across different viral batches to assess differences in expression ratios of the viral proteins and, also, to monitor for maturation of the GP complex (GPC) which occurs via cleavage into the GP1 and GP2 subunits. Our results demonstrate that batch to batch variability was minimal at the protein level. Furthermore, we demonstrate that, as expected, the GP glycoprotein was efficiently processed over time into its individual subunits with ratios indicating little or no shedding from the virus particle. This UPLC-MRM assay has proven to be a powerful method for measuring viral vector proteins which in turn can provide better understanding of complementary data monitoring infectivity, potency, and stability of different viral batches (30, 31).

We then further refined the assay to include quantification of the full-length GPC before processing to GP1 and GP2 subunits. We and others have observed evidence for GPC processing employing semi-quantitative methods like Western Blot (**Fig. S1**) (32, 33). The GPC undergoes proteolytic processing by site 1 protease (S1P) yielding two subunits, the 44-kDa peripheral GP1 and the 35-kDa transmembrane GP2. The prefusion form of GP1-GP2 exists as a protomer (34, 35) with the outer GP1 responsible for receptor engagement with its target receptor, α-dystroglycan (α-DG). The receptor binding results in entry of the virus into the cells via endocytosis. Subsequently, the acidic pH in the endosome triggers the dissociation of GP1 from GP2 and conformational changes in GP2 drive the fusion of the virus to host membranes. By independently measuring GP1 and GP2 concentrations, the GP1/GP2 ratio can provide an assessment of the stoichiometry of the envelope protein complex which is known to be important for infectivity (32, 36). The UPLC-MRM method developed in this study offers a new and complementary approach to provide specific and quantitative insight not only into the concentrations of GP1 and GP2 on the surface of the viral particles but also for any remaining unprocessed GPC. Solely monitoring GP1 and GP2 levels will not identify whether the GP1 and GP2 peptide surrogates are being detected from the processed or full-length unprocessed GPC. Consequently, our final method contained an MRM transition for a single tryptic peptide surrogate that is specific for the full-length unprocessed GPC in the viral samples. Using this approach, the temporal kinetics of GPC expression and processing could be evaluated in the production cells by simultaneously monitoring the changes between the three GP components (unprocessed full-length GPC, and the processed products GP1 and GP2) which provided insight into the viral lifecycle.

## RESULTS

### VSV-GP proteins N, P, M, GP1, GP2, and L identified by surrogate peptides

It is essential for a robust MRM assay to have the correct protein sequences and to identify suitable and specific surrogate peptides corresponding to the proteins of interest. Thus, *in-silico* sequences were first obtained from the National Center for Biotechnology Information (NCBI) database with all protein sequences from VSV except for the G-protein since the VSV used in our studies had been pseudotyped with glycoprotein from LCMV (23). To confirm these sequences were present in VSV-GP, we first carried out peptide mapping experiments using a Q-TOF mass spectrometer (Synapt G2, Waters, Milford, MA). Once the sequences were confirmed, we then proceeded to use Skyline software (University of Washington) to generate transition lists to determine potential surrogate peptides for each viral protein (37). Skyline generated virtual tryptic digests of the sequences and created a list of transitions corresponding to the individual tryptic fragments (37). The software allows the transitions and method parameters to be customized for different instrument types including our Sciex triple stage quadrupole instrument. Multiply charged precursor ions were chosen with fragment ions having *m/z* values higher than the precursor ion to improve selectivity (38). Since we intended to use C-terminally labelled internal peptide standards, y fragment ions were specifically selected for Skyline generation of transitions (**Table 1**) (14, 39, 40). The sequences generated from Skyline were exported to the Analyst Software for use with a Sciex 6500 triple quadrupole mass spectrometer.

**Table 1:**
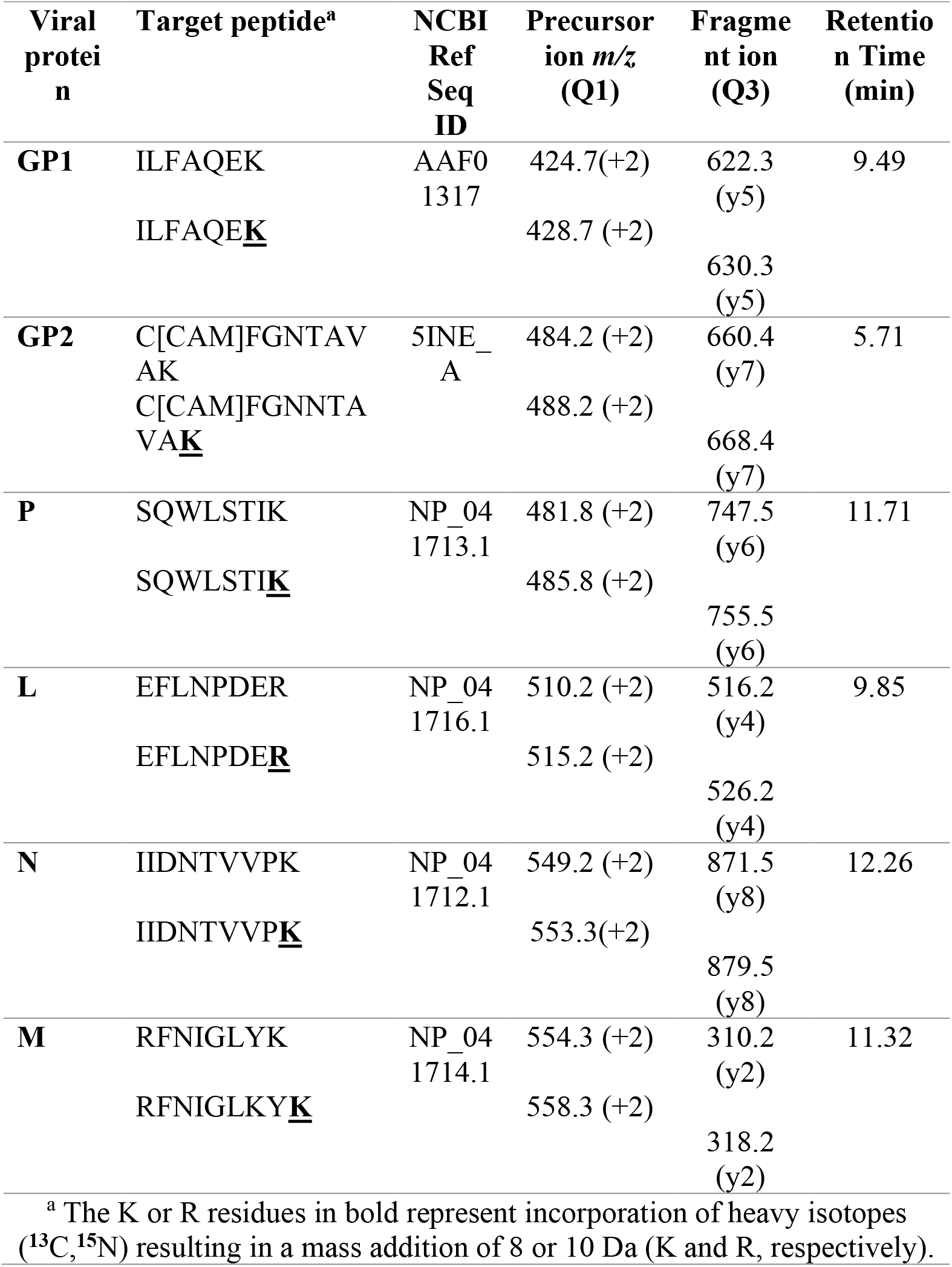
Mass transitions and MRM parameters of light and heavy labeled surrogate peptides for the VSV-GP proteins in the study.

Next, we empirically tested all the transitions to determine the optimum surrogate peptide transitions using purified virus (Batch 1) that had been subjected to trypsin digestion. The resultant UPLC-MRM chromatograms and a list of the most intense transitions can be found in **Fig. S2** and **Supplementary XS1**, respectively. More than two transitions corresponding to a particular precursor ion at the same retention time provided a first line confidence check for the specificity of the transitions (41). Specificity of potential surrogate peptides for the intended viral protein was then further confirmed by carrying out a NCBI Protein BLAST search to ensure peptide uniqueness. Peptides confirmed to be specific to VSV-GP and having transitions providing the maximum possible peak area were then selected for surrogate peptide synthesis (**Table 1**).

We next performed the same process to identify surrogate peptides that were specific to the processed GPC. Identification of a surrogate peptide for full-length GPC was more complex and is discussed in the next section. Once all the optimum surrogate peptides had been identified, both labeled and unlabeled forms of the surrogate peptides were synthesized for use as standards.

### Identification and selection of surrogate peptides for GPC extends the potential of the assay for monitoring full-length GPC

The surrogate peptide for full-length GPC was selected as the sequence spanning both GP1 and GP2, **Fig. S4** (33). Unlike for GP1, GP2, and the other viral peptides, the initial analysis of Batch 1 used for method development did not identify any detectable transitions for a full-length surrogate of GPC (**Table 3**) suggesting efficient GPC processing in the purified virus sample. However, we were ultimately able to detect two transitions specific for the full-length GPC (from the surrogate peptide having the sequence LAGTFTWTLSDSSGVENPGGYCLTK) after using a larger starting volume of purified virus which allowed us to concentrate the digest approximately 10-fold using a speed-vac concentrator (**Fig. S4**). Analysis of this concentrated sample by our UPLC-MRM method allowed detection of two transitions at exactly the same retention time (20.8 min) which demonstrated these two transitions for this surrogate peptide could be used to monitor unprocessed full-length GPC **(Table 2**). Targeted analysis of both this GPC surrogate peptide and the GP1 and GP2 surrogates extends the potential of the assay, enabling the assessment of GPC processing which is discussed later.

**Table 2:**
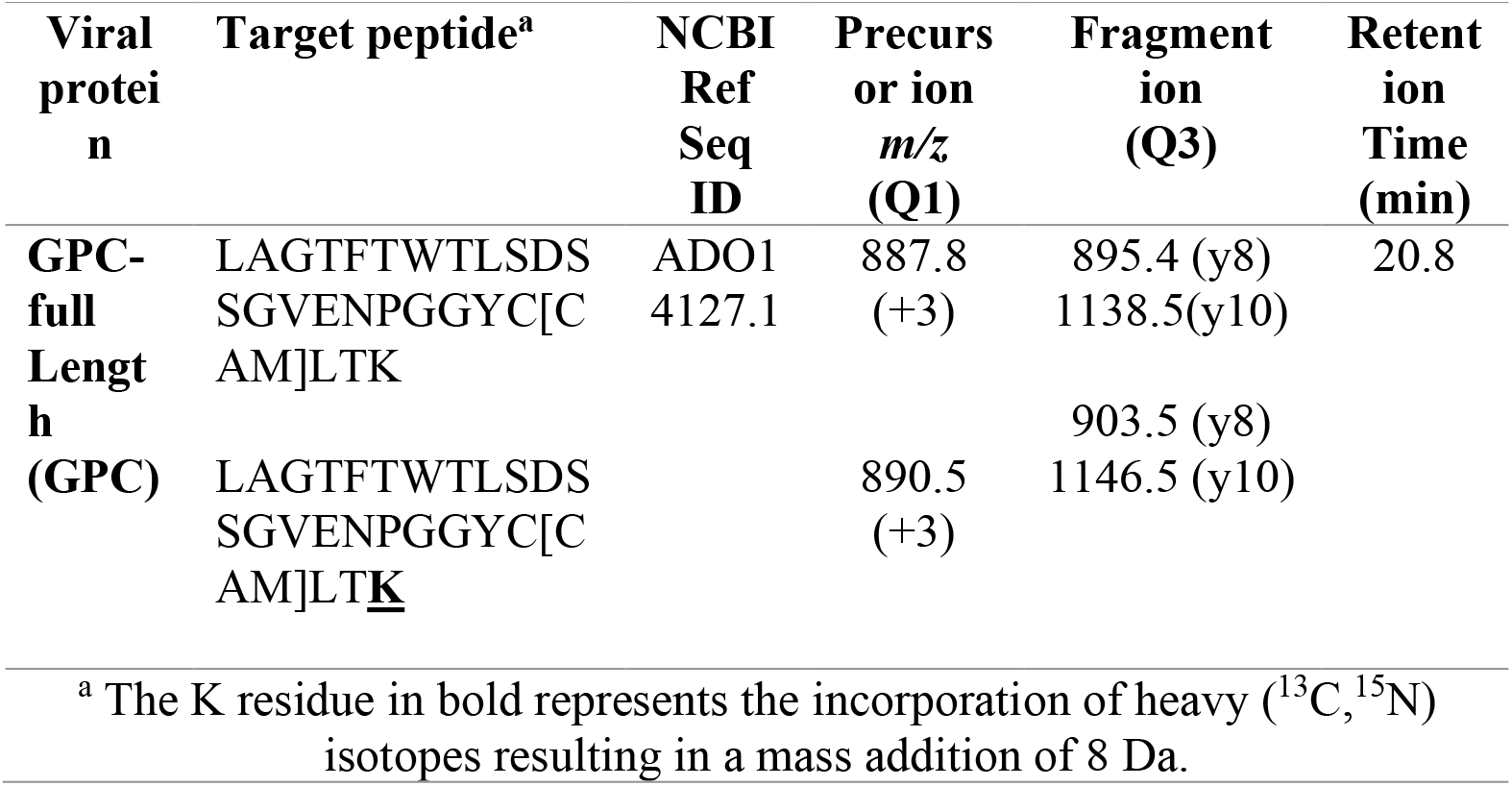
Mass transitions and MRM parameters of light and heavy labeled surrogate peptides for full-length GPC.

**Table 3:** Concentration of surrogate peptides and corresponding VSV-GP full-length protein concentrations in Batch 1a.

| **Viral Protein** | **Concentration of peptide (ng/mL)<sup>b</sup>** | **Monoisotopic Mass (Da)** | **Concentration of protein (μM)<sup>c</sup>** | **Coefficient of Variation (%)<sup>d</sup>** | **Lower limit of quantification (LLOQ)<sup>e</sup> (μM)** |
| --- | --- | --- | --- | --- | --- |
| **GP1** | 370±5 | 847.48 | 0.44±0.02 | 5 | 0.002 |
| **GP2** | 648±10 | 968.04 | 0.55±0.05 | 9 | 0.008 |
| **P** | 317±2 | 961.52 | 0.33±0.04 | 12 | 0.002 |
| **L** | 108±5 | 1018.47 | 0.07±0.01 | 12 | 0.001 |
| **N** | 1069±22 | 1097.65 | 1.42±0.2 | 15 | 0.002 |
| **M** | 3067±35 | 1106.62 | 2.8±0.2 | 14 | 0.012 |
| **GPC<sup>f</sup>** | ND | 2603.22 | ND | ND | 0.003 |

<sup>a</sup> Batch 1 was used to identify and optimize the MRM surrogate peptide transitions.
<sup>b</sup> The absolute concentrations of the surrogate peptides (Table 1) were calculated using MultiQuant 3.0 software. Values are the mean and standard deviation from three independent replicates (duplicate injections of each replicate).
<sup>c</sup> The molar protein concentrations were determined by converting surrogate peptide concentrations (ng/mL) to molar concentrations (μM).
<sup>d</sup> The coefficient of variation is reported for triplicates.
<sup>e</sup> LLOQ was determined by injecting the lowest concentration of each dilution series for individual peptides and using the ICH guidance LLOQ = 10 \* SD/S, where SD is the standard deviation of the six injections and S is the slope of the standard curve generated for the peptide (**45**).
<sup>f</sup> Although GPC transitions (Table 2) were determined from Batch 1, absolute concentration measurements could not be determined (ND) due to very low GPC abundance.

### Quantifying the major viral proteins with high precision and accuracy

The relative amounts of the viral proteins in native VSV have previously been reported by other groups using techniques like western-blot and transmission electron microscopy (27, 30, 42–44). We used this information together with our own initial method development data to design the standard concentration dilutions for each peptide standard such that all sample measurements were well within the linear range (**Fig. 1**, **Table 3**). For example, the red circles in **Fig. 1** denote the positions of the Batch 1 sample peptide concentrations on the linear regression line for each surrogate peptide.

**Fig. 1:**
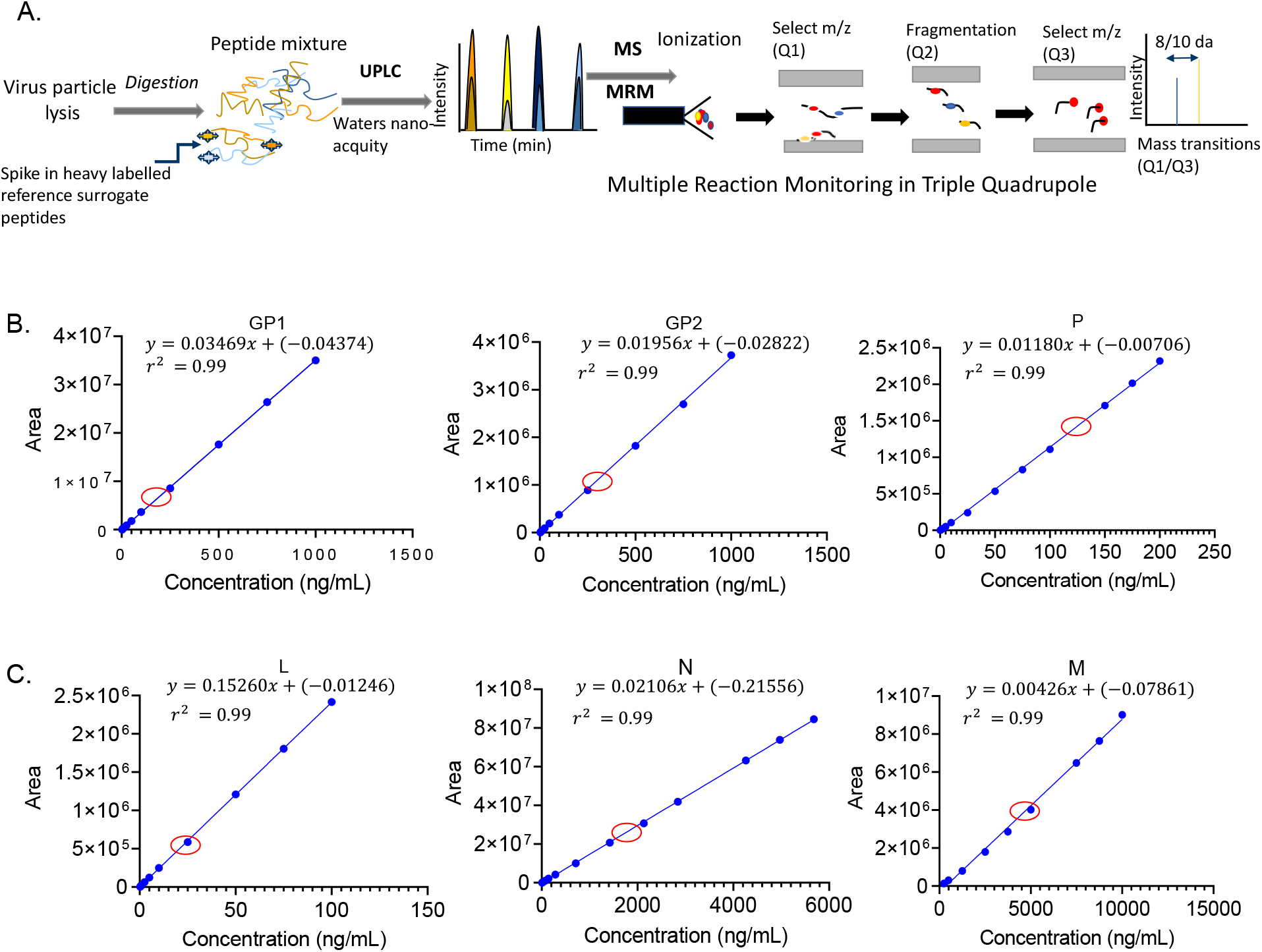
Schematic representation of the method and surrogate peptide standard curves. **A.** The method used for peptide identification involves lysing the viral particles followed by trypsinization using an Endo-Lys-C/trypsin protease cocktail. The resulting tryptic peptides are separated using UPLC and detected via MRM on a Sciex 6500 triple stage quadrupole instrument. This enables absolute quantification of the proteins corresponding to the surrogate peptides. Use of MRM detection makes this method very specific and selective for the viral proteins of interest. **B, C** Standard curves using unlabeled surrogate peptides for GP1, GP2, P (top Panel **B**) and L, N and M (bottom panel **C**) are shown. The standard concentration ranges for the peptides were adjusted empirically based on the actual concentration range of the proteins present in the virus, GP1 (0.5-200 ng/mL), GP2 (0.5-200 ng/mL), P (0.5-100 ng/mL), L (0.5-100 ng/mL), N (7.1-5680 ng/mL) and M (25-5000 ng/mL). The red circles on the graph denote the Batch 1 sample peptide concentrations. Linear regression using 1/x weighting was done with MultiQuant 3.0 software which calculated the equation and r^2^ values for each plot. The lowest concentration of each peptide was injected 6 times to obtain the lower limit of quantification (LLOQ, **Table 3**). Each linear regression (standard curve) was generated from three independent runs with duplicate injections thus creating an average of six data points for each concentration. The error bars represent the standard deviation of those six data points for each concentration.

Each standard curve was derived from three technical replicates (each injected twice) with calculated r^2^ values ≥0.98 for all standard curves.

A major strength of using mass spectrometry detection is that precursor ions from heavy labeled peptides have equivalent retention time (and fragmentation pattern) to the non-labeled surrogate peptide yet have a higher mass (8 or 10 Da) due to the presence of the label **(Fig. S3, Table 2).** This feature allows unlabeled and labeled peptides with the same sequence to be separated by Q1 in the mass spectrometer despite having essentially identical retention times (39). In this method, the same exact amount of synthesized heavy labeled peptide is added to both external standards and samples. While the amount of native (light) peptide changes depending on the external standard dilution preparation the amount of heavy peptide is kept the same. Since the light and heavy peptides behave identically but can be separated by a mass spectrometer based on mass, this provides a very accurate and precise method for absolute concentration determination that is not affected by variables such as, for example, MS signal response due to slight changes in the ion source electrospray or small differences in injection volume. Therefore, the x-axis in the standard curve graphs is a concentration ratio rather than concentration since the peak area is normalized to the peak area of the labeled reference peptide (46) as shown in **Fig. S6** with this data also demonstrating the accuracy and precision of the method.

The development of the UPLC-MRM method for absolute quantification with an isotopically labeled internal standard was based on the viral protein levels in VSV-GP Batch 1. The absolute quantification of viral proteins in Batch 1 was done by spiking in the same amount of heavy internal standard 50 ng/mL (for GP1, GP2, P, L and N surrogates) or 250 ng/mL (for the M surrogate). As the protein M was empirically observed to be present in much higher amounts, a 5-fold higher concentration of M internal standard was used. Each sample was examined as a technical triplicate run on the same day with three independent biological measurements performed over several days. Therefore, we were able to report both intraday variability (from the technical replicates) and interday variability (from the biological replicates) **(Fig. 2**, **Table 3)**. The Coefficient of Variation (CV) for all concentration measurements was ≤15 % demonstrating the excellent reproducibility of the assay **(Table 3)**. Furthermore, the relative values of each of the structural proteins for VSV-GP were comparable to that reported in the literature with M and N having the highest amounts and P and L having the lowest amounts (27, 30, 47). The absolute concentrations of each of the proteins were calculated and expressed as molar amounts (µM) **(Table 3**, **Fig. 2)** but were also calculated as relative concentrations [relative to M] since this is the most abundant viral protein in VSV-GP **(Fig. 2**, **Table 5).**

**Fig. 2:**
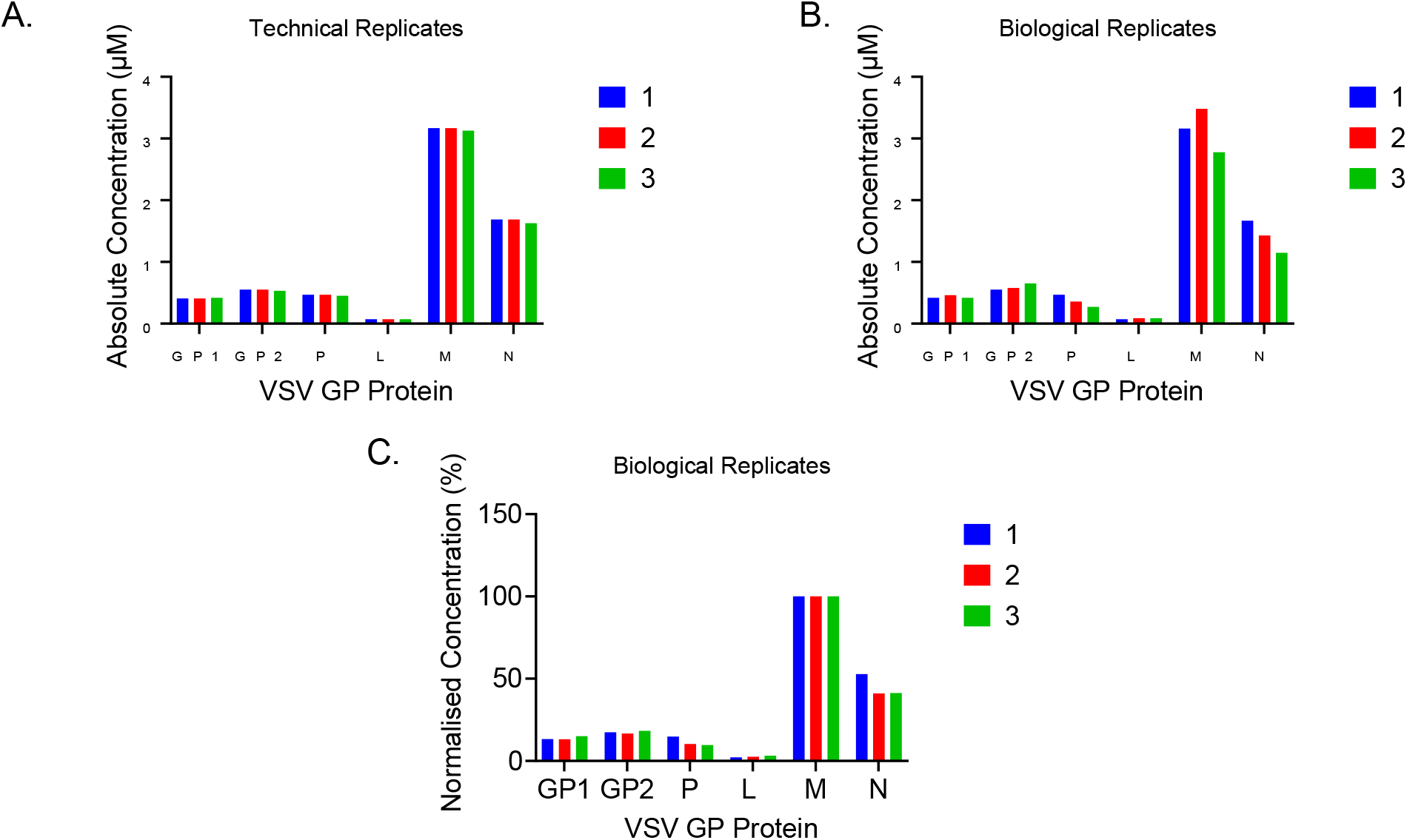
Concentrations of VSV-GP proteins in Batch 1. A) Histograms of three technical replicates of the same sample demonstrating intra-day variability between runs. The absolute micromolar protein concentrations were calculated from the concentration of the peptide reported by MultiQuant 3.0 in ng/mL and the known molecular weight of the peptides. B) Histograms showing absolute concentrations of the VSV-GP proteins with three independent biological samples, performed on three independent days, showing inter-day variability. Mean concentrations with corresponding standard deviations are reported in **Table 3**. C). The concentrations of the VSV-GP proteins normalized to the concentration of M (the most abundant protein). The normalization has been done on the biological replicate samples for which absolute values are plotted in (B).

Determination of the stability of the peptide surrogate standards is required to identify appropriate storage conditions to provide accurate and reliable measurements (48). To characterize in-use stability, UPLC-MRM analysis was also carried out on standard solutions (dilutions) kept in storage at 4 °C for 7 days (**Fig. S5**). We observed that for all peptide curves the slope changed after 7 days compared to measurements made for standard dilutions prepared freshly from stock. The most extreme deviation was observed for the P protein standard peptide which indicated this peptide was unstable and had undergone significant degradation within our experimental time span. To determine long-term storage stability, the stock solutions were stored at −25 °C and standard curve calculations were performed after one month (two freeze-thaw cycles). The slope and the r^2^ values were reproducible when compared to the freshly prepared standards (data not shown). We therefore concluded that the stock solution of peptides could tolerate storage at −25 °C for one month without undergoing degradation and could be used for standard curve generation. However, we found that prolonged storage at 4 °C was not viable.

### Comparison between different viral batches show consistency

We then used our optimized method to compare viral protein abundances (and confirm GPC processing) in several purified virus batches generated during process development. For this, we selected 7 additional batches (Batch 2-8) and compared the protein concentrations of each batch relative to Batch 1 (**Fig. 3**, **Table 4**). Batch 1 showed a higher absolute amount of each GP1, GP2, P, L, N and M proteins with the total viral protein amounts of each batch tracking well with the genome content established separately by real-time quantitative reverse transcription PCR (qRT-PCR) and the infectious viral titer which was estimated by calculating average 50% Tissue Culture Infectious Dose (TCID_50_) (**Table S2**). For each batch it was observed that the M protein was the most abundant, followed by protein N with P and L present at the lowest levels, in accordance with findings by Thomas et al (42). Since the total viral protein concentrations for each batch correlated with measured genome content, we also normalized the protein concentrations in the same batch relative to the most abundant protein M to better understand the protein concentration distributions within batches. After normalization the viral protein ratios were nearly identical in each batch as indicated in **Fig. 3B**, **Table 5**. The GPC concentrations could not be determined (as would be expected if GPC is mostly if not fully processed in the purified batches) which led us to conclude the observed GP1 and GP2 concentrations were measurements of GP1 and GP2 subunit proteins after GPC processing.

**Fig. 3:**
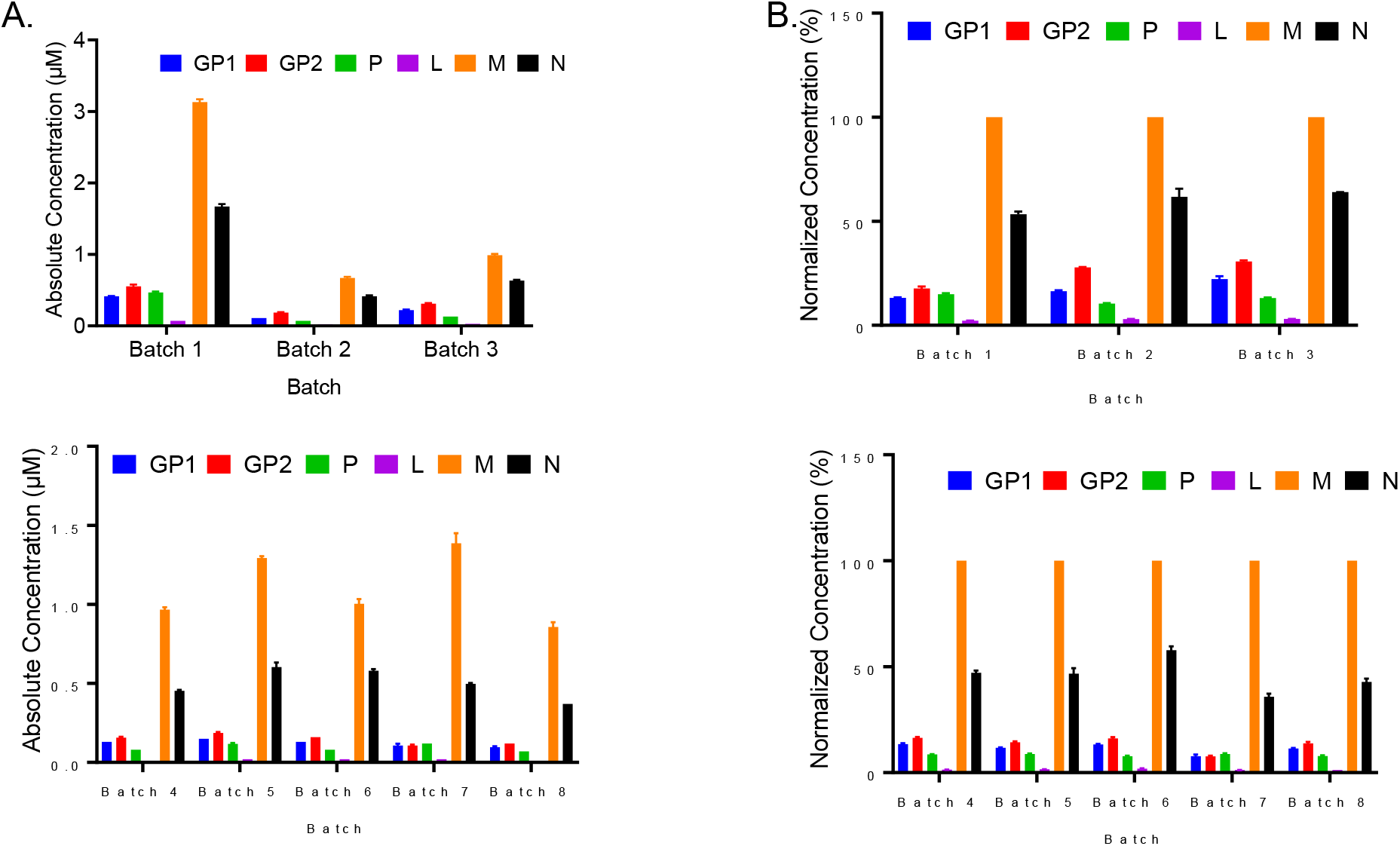
Comparison of absolute and normalized concentrations of the VSV-GP proteins, between different viral batches: A) (Left Panel) Absolute concentrations (µM) of VSV-GP proteins from different developmental batches (1–8). B). (Right Panel) shows the same data but with concentrations normalized to the most abundant protein M. In all cases the plotted histograms are the mean of two samples from each batch with each sample run in triplicates, error bars representing standard deviation (**Table 4**).

**Table 4:** VSV-GP absolute protein concentrations for different purified viral batches.

|  |  |  |  |  |  |  |
| --- | --- | --- | --- | --- | --- | --- |
| **Viral Batch** | **Protein Concentration (μM) <sup>a</sup>** |  |  |  |  |  |
|  | **GP1** | **GP2** | **P** | **L** | **N** | **M** |
| **Batch 1<sup>b</sup>** | 0.42±0.01 | 0.55±0.01 | 0.47±0.01 | 0.07±0.00 | 1.67±0.03 | 3.16±0.02 |
| **Batch 2** | 0.21±0.01 | 0.24±0.01 | 0.09±0.01 | 0.03±0.01 | 0.47±0.02 | 0.95±0.01 |
| **Batch 3** | 0.22±0.01 | 0.31±0.02 | 0.13±0.02 | 0.03±0.01 | 0.63±0.01 | 0.99±0.01 |
| **Batch 4** | 0.13±0.01 | 0.15±0.02 | 0.08±0.01 | 0.01±0.00 | 0.45±0.02 | 0.98±0.01 |
| **Batch 5** | 0.15±0.02 | 0.18±0.01 | 0.12±0.01 | 0.02±0.01 | 0.60±0.03 | 1.29±0.02 |
| **Batch 6** | 0.13±0.01 | 0.16±0.01 | 0.08±0.01 | 0.02±0.01 | 0.58±0.01 | 1.00±0.01 |
| **Batch 7** | 0.11±0.01 | 0.12±0.01 | 0.12±0.01 | 0.01±0.01 | 0.50±0.02 | 1.39±0.01 |
| **Batch 8** | 0.11±0.01 | 0.12±0.01 | 0.07±0.01 | 0.01±0.00 | 0.37±0.01 | 0.89±0.01 |

<sup>a</sup> Each batch was sampled twice with each sample run in triplicate. The values reported here are average of the (6) determinations showing standard deviation. <sup>b</sup> Batch 1 is the batch used for development of our UPLC-MRM assay
<sup>b</sup> Batch 1 is the batch used for development of our UPLC-MRM assay

**Table 5:** VSV-GP protein concentrations of different purified viral batches normalized to M.

|  |  |  |  |  |  |  |
| --- | --- | --- | --- | --- | --- | --- |
| **Viral Batch** | **(%)<sup>a</sup>** |  |  |  |  |  |
|  | **GP1** | **GP2** | **P** | **L** | **N** | **M** |
| **Batch 1<sup>b</sup>** | 13.2±0.2 | 17.7±0.8 | 14.9±0.2 | 2.5±0.1 | 53.4±1.1 | 100 |
| **Batch 2** | 16.4±0.3 | 25.2±0.01 | 10.5±0.1 | 3.0±0.1 | 61.7±2.3 | 100 |
| **Batch 3** | 22.3±1.11 | 30.6±0.5 | 13.2±0.2 | 3.03±0.1 | 63.9±0.1 | 100 |
| **Batch 4** | 13.4±0.4 | 16.3±0.4 | 8.6±0.0.1 | 1.4±0.1 | 47.1±0.8 | 100 |
| **Batch 5** | 12.10±0.2 | 14.0±0.4 | 8.8±0.1 | 1.6±0.1 | 46.8±1.2 | 100 |
| **Batch 6** | 13.2±0.2 | 16.1±0.01 | 7.8±0.1 | 1.8±0.2 | 56.7±1.5 | 100 |
| **Batch 7** | 7.7±0.6 | 7.6±0.3 | 8.8±0.3 | 1.2±0.1 | 36.0±0.2 | 100 |
| **Batch 8** | 11.3±0.2 | 13.8±0.3 | 7.8±0.3 | 1.3±0.1 | 42.8±1.1 | 100 |

<sup>a</sup> Each batch was sampled twice with each sample run in triplicate. The values reported here are average of the (6) determinations showing standard deviation from the mean.
<sup>b</sup> Batch 1 is the batch used for development of our UPLC-MRM assay

Processing of full-length GPC into GP1 and GP2 is a prerequisite for membrane transport of the virus and hence infectivity (49) (32). In addition, the outer GP2 protein associates with the membrane bound GP1 but not covalently (34). For this reason, it was important to measure the GP2/GP1 ratio, which can inform on both GPC processing and possible “stripping” of GP1 from the viral particle surface - it has been shown earlier that during the infection process GP1 may get dissociated from GP2 resulting in a slightly higher GP2 content (34, 50). Previous studies have also revealed the presence of unprocessed GPC in certain viral particles with both fully and partially processed GPC determined by semi-quantitative methods such as western blot assays (49, 51). Building on the previous knowledge, we now have additional quantitative confidence with the UPLC-MRM approach since absolute concentrations of unprocessed full-length GPC as well as the post-processed subunits GP1 and GP2 can be monitored simultaneously. Our results suggest that the absolute quantities of GP1 and GP2 in all the viral batches were comparable, resulting in a GP2/GP1 ratio at or near 1.0 for most batches (**Table 4**). Although other studies have shown that GP1 may get dissociated from GP2 during the infection process resulting in a slightly higher GP2 content (50)in our VSV-GP data the ratio of GP2 to GP1 in all tested batches remains very close to 1.0, suggesting not just complete processing of GPC, (32, 50, 52) but also no evidence for any significant GP1 “stripping” off the surface of the viral particles.

The detection of MRM transitions specific to unprocessed GPC allowed us to incorporate GPC specific surrogate heavy and light peptide standards into our assay. Empirically, it was observed that at concentrations lower than 5 ng/mL the unlabeled GPC peptide surrogate showed poor signal response which was not altogether surprising given the longer peptide length (27 amino acids) (53). Consequently, higher concentrations for the GPC peptide surrogate were used compared to the other VSV-GP surrogate peptides for generation of the GPC standard curve. A good linear response (r^2^ ≥ 0.98) could be obtained within the concentration range of 15 ng/mL-1200 ng/mL with a LLOQ of 0.003 nM providing excellent sensitivity for GPC (**Table 3, Fig. S7**). After using a larger sample volume and concentrating via a SpeedVac, the levels of GPC surrogate peptide in Batch 1 were ultimately detectable but with very low signal to noise which did not allow confident calculation of GPC concentration.

### GPC processing can be tracked in VSV-GP infected cells over time with the UPLC-MRM assay

The quantitative measurements of the viral proteins (GP1, GP2, L, M, N, and full-length GPC) discussed up until now were all obtained from purified virus batches. We next wanted to examine how the protein levels might change over time. We were specifically curious to see if we could capture the progress of GPC processing by exploiting the sensitivity and selectivity of this assay for independently tracking GP1, GP2, and GPC. For this, we infected HEK293-F (production cells) with VSV-GP at multiplicity of infection (MOI) of 0.1 and collected the cells at 5 min, 3 h and 30 h time points for lysis and protein quantification by UPLC-MRM.

We observed a decreasing trend in GPC peak intensities in the 5 min, 3 h and 30 h time point samples. The GPC peak in the 5 min sample was sufficiently abundant (within the range of the standard curve) for the absolute concentration to be easily calculated (0.02±0.01 µM). However, no peaks were detected for GP1 and GP2 (**Fig. 4**, **Table 5**). This supported the hypothesis that GPC-processing to GP1 and GP2 is incomplete at the initial stage of infection. Although the 3 h sample showed nearly a 2-fold lower level of GPC (0.01 µM) compared to the sample taken at 5 min, GP1 and GP2 remained undetectable. On the other hand, the 30 h time point sample had no detectable GPC but measurable levels of GP1 and GP2 (both quantified at 0.03 µM). The concentrations of GP1 and GP2 measured in these samples were significantly lower than in the other viral batches we analyzed but this was expected since the latter were purified virus batches (final drug product) and, hence, much more concentrated. Similarly, the remaining VSV-GP proteins, while detected in the 5 min and 3h time point samples, also had lower levels compared to concentrations that would be expected in the drug product. All of the viral proteins (with the exception of GPC) were easily detected in the 30h time point sample. The other VSV-GP proteins (P, M, N and L) also showed an increasing trend of concentration in the time -point samples, with L being detectable and quantifiable in the 30 h sample. This provides further support for our hypothesis that this UPLC-MRM method can track viral maturation by analysis of time-point samples (**Fig. S7**). Mock cells (having no viral infection) were used as controls with no surrogate peptides detected for any of the VSV-GP proteins including GPC for all time points (**Table 6**). In addition, the complete viral life cycle was confirmed by TCID_50_ assay, which was carried out on supernatants collected at same time points (5 min, 3 h and 30 h) to quantify infectious virus progeny release from cells (**Fig. S7**). In summary, we were able to observe expected differences in GPC levels between the time point samples that provided support for our hypothesis and demonstrated that our assay could indeed detect unprocessed GPC along with a concomitant decrease in GP1 and GP2 levels.

**Fig 4:**
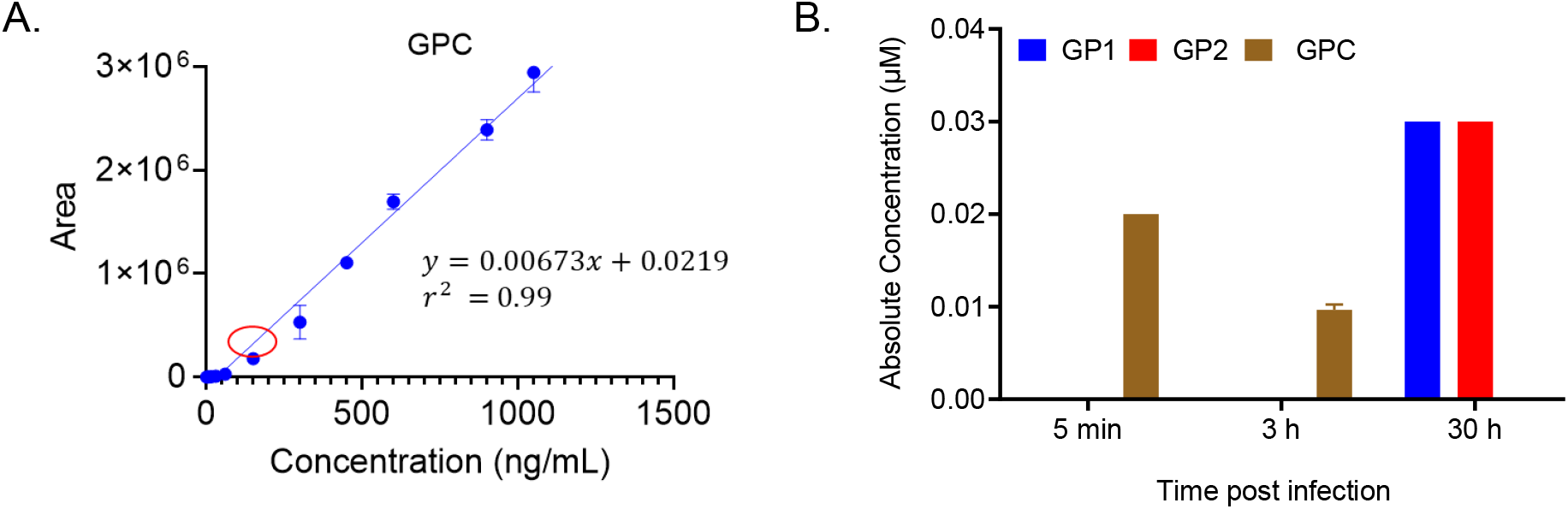
Full-length GPC detection in time point samples of HEK-293 cells infected with VSV-GP: A) Standard curve with unlabeled surrogate peptide for full-length GPC (1.-1200 ng/mL), the red circle showing concentration location for the earliest time point sample (5 min) for which full-length GPC could be quantified. B) Concentrations of GP1, GP2 and full-length GPC in time point cell pellets of HEK-293F cells infected with VSV-GP. The plotted histograms are the mean of two samples from each batch with each sample run in triplicate with error bars representing standard deviation. C) Time course D) Also detectable in samples after DSP

**Table 6:** Protein concentrations for VSV-GP infected time point samples and uninfected samples.

| Table 6: Protein concentrations for VSV-GP infected time point samples and uninfected samples <sup>a</sup> |  |  |  |  |  |  |  |
| --- | --- | --- | --- | --- | --- | --- | --- |
| Viral Batch | Protein Concentration (μM) |  |  |  |  |  |  |
|  | GP1 | GP2 | GPC | L | N | M | P |
| 5 min | ND | ND | 0.02±0.01 | ND | 0.02±0.01 | 0.02±0.00 | 0.01±0.02 |
| 3 h | ND | ND | 0.01±0.00 | ND | 0.01±0.01 | 0.02±0.01 | 0.01±0.02 |
| 30 h | 0.03±0.01 | 0.03±0.01 | ND | 0.01±0.00 | 0.19±0.02 | 0.34±0.02 | 0.07±0.03 |
| Mock infection 5 min, 3h, 30 h | ND | ND | ND | ND | ND | ND | ND |

<sup>a</sup> The samples were run in technical triplicates. The values reported here are average of all runs with standard deviation from the mean. ND represents Not Detected.

## DISCUSSION AND CONCLUSION

The rapidly developing space of oncolytic virus therapy exemplified by VSV-GP necessitates the use of novel analytical tools for characterization that are also accurate, precise, and sensitive. In the present work we have harnessed the analytical power of UPLC-MRM using both isotopically labeled and non-labeled surrogate peptide standards to develop an assay for absolute quantification of the structural proteins encoded by VSV-GP (GP1, GP2, P, L, N and M). The mature assay provided linear standard curves with nanomolar lower limits of quantification for all proteins with excellent reproducibility and coefficients of variation ≤15%. The use of known amounts of heavy labeled surrogate peptide added to each standard and sample increased the accuracy and precision and thus, the confidence of the quantification.

We analyzed the absolute concentrations of the VSV-GP proteins in different purified viral batches that informed us on the similarities and differences between these batches. In summary, our results demonstrated a consistent trend in the protein content, with M and N being the two most abundant proteins and P and L being the least abundant(42, 54–57). The differences in the absolute quantity of VSV-GP proteins between the batches may reflect slight differences in the production process (which is beyond the scope of this study).

The VSV-GP viral envelope protein is important for virus infectivity. We utilized separate surrogate peptide standards for specific quantification of the two subunits GP1 and GP2 that result after complete processing of the LCMV-GP protein (GPC). We also identified a third tryptic peptide that could be used as a surrogate for GPC since it could only exist if GPC had not been fully processed to GP1 and GP2. If unprocessed GPC was not detected (as was the case in all the virus batch samples tested) we assumed that any GP1 and GP2 detected arose primarily from the processed subunits. Monitoring full-length GPC together with the processed GP1 and GP2 subunits also allowed us to track GPC processing. With Batch 1 used for developing the assay, the GPC peptide was only detected after using a larger starting volume and concentration but the signal due to GPC was still below the limit of quantification of the assay. For the other batches (Batch 2-8) no GPC surrogate peptide was detected. Therefore, we concluded that in the purified viral batches, the GPC processing is essentially complete with all GPC converted to GP2 and GP1. We further tested this hypothesis by generating samples expected to contain GPC that had not been completely processed. Specifically, we infected HEK-293F cells with VSV-GP and took time point samples (5 min, 3 h and 30 h) with the expectation that the earlier time points would have measurable levels of unprocessed GPC. Our results show a clear distinction in the levels of GPC, GP2 and GP1 detected over time, with GPC being most abundant in the 5 min and to a lesser extent in the 3 h time point samples and with both 5 min and 3 h samples containing no detectable GP2 and GP1. On the other hand, GP2 and GP1 were present in quantifiable amounts while GPC was not detected in the 30 h time point sample. These results demonstrate that the UPLC-MRM assay we developed can provide important information on viral maturation as well as a snapshot of GPC processing by simultaneously measuring the levels of GPC, GP1, and GP2. In addition to tracking GPC processing, the ability to analyze GP1 and GP2 separately enabled us to determine the GP2/GP1 ratio which allows monitoring of any potential “shedding” of the external non-membrane bound GP1 subunit of the GP2/GP1 complex with higher GP2/GP1 ratios potentially an indication of GP1 dissociating from GP2 (35, 52). As expected, the GP2/GP1 ratios were generally uniform in our purified virus batches with little or no observed “stripping” of the outer non-membrane bound GP1.

In the present work, we have established the UPLC-MRM assay as a useful tool for multi-plex absolute quantification of viral proteins in VSV-GP batches. The assay has the advantage of being a multiplexed, selective, accurate and precise, highly sensitive, reproducible, and absolute quantitative method for analysis of viral proteins for oncolytic viruses and potentially other viral vectors. Although the method presented here has been used for the absolute quantification of viral proteins from pseudotyped VSV-GP, it has the flexibility to be adapted for quantifying and characterizing other proteins of interest such as host cell proteins and therapeutic cargo proteins. Finally, our assay can be extended as a bioanalytical tool to other viral therapy platforms. (word count 3336)

## MATERIALS AND METHODS

### Chemical Reagents and Materials

All experiments were carried out using Optima^®^ LC/MS grade (Fisher Scientific, NY) water, organic solvents and acids. Temperature dependent incubations for the processing of viral samples and trypsinization were done using an Eppendorf Thermomixer. RapiGest for viral lysis was purchased from Waters Inc, (Milford, MA). The synthesis of the surrogate peptides both labeled and unlabeled was done by GenScript (Piscataway, NJ) and provided in lyophilized form.

### Viral batch production

VSV-GP virus batches were produced in HEK-293F cells in either shaker flask or bioreactor formats (depending on batch ID). After infecting cells at an MOI of 0.0005, virus was purified from clarified harvest via centrifugation and/or chromatographic methods similar to previous reports (58, 59). The downstream processes employed were slightly unique for each batch (for example the use of different purification columns or formulations).

### Viral protein preparation

An aliquot of purified virus obtained from Development batches was concentrated in a 50 mM ammonium bicarbonate solution by centrifugation at 14,800xg for 2 h. The supernatant was then carefully removed, with the concentrated virus resuspended in 0.1 % RapiGest diluted in 50 mM ammonium bicarbonate (pH 7.8) (60). For lysis of the virions, the suspension was sonicated for 15 min and then heated to 90 °C for 1 h with 300 rpm shaking before a further 15 min sonication. The samples were reduced by the addition of tris (2-carboxyethyl) phosphine (TCEP) to a final concentration of 20 mM and incubated at 60 °C with 600 rpm shaking for 1 h. The samples were then cooled to room temperature and alkylated by the addition of iodoacetamide to a final concentration of 40 mM and incubated at 37 °C with 600 rpm shaking for 30 min in the dark. A mixture of trypsin and Endo-Lys-C (Trypsin/Lys-C Mix, Mass Spec Grade, Promega) was used for digestion of the sample. Trypsin/LysC was reconstituted in 50 mM ammonium bicarbonate to a concentration of 1 µg/ml. Samples were digested by addition of 4 µL trypsin, followed by incubation at 37 °C overnight. The digestion was terminated by acidification (addition of trifluoroacetic acid to a final concentration of 0.5% v/v and incubation at 37 °C for 1 h with 600 rpm shaking) followed by centrifugation (13,000xg for 10 min). Prepared samples not run immediately by UPLC-MRM were stored at −20 °C.

### Calibration curve with light peptide standards

The unlabeled (light) surrogate peptides corresponding to the respective viral proteins GP1, GP2, P, L, N and M were synthesized by GenScript to more than 98% purity except for the surrogate peptide for full-length GPC which could only be purified to 66%. All synthetic peptides were supplied in a lyophilized form. Initial stock solutions were prepared to a concentration of 1 mg/mL by weighing and reconstituting the lyophilized peptides in LC-MS grade water having 0.1 % trifluoroacetic acid. A standard cocktail solution with all six peptides GP1, GP2, P, L, N and M was prepared with concentrations 100, 100, 10, 10, 280 and 500 ng/µL, respectively. Serial dilutions of this cocktail stock solution were then prepared to cover the expected concentration range of the peptides. For GP1 and GP2 peptides, this dilution range corresponded to 2.5 – 1000 ng/mL, 0.5-200 ng/mL for P, 0.5-100 ng/mL for L, 7.1-5680 ng/mL for N and 25-5000 ng/mL for M. The GPC stock solution was prepared separately at a concentration of 60 ng/µL. The stock solution of GPC was serially diluted to cover the concentration range 1.5-1200 ng/mL. The standard solutions were prepared in approximately 2-fold increments covering the entire concentration range. Standards were injected from lower concentration to higher concentration and each injection was done at least twice. For all analyses done in this work, blank injections having only LC-MS grade water containing 0.1 % trifluoroacetic acid were used liberally. For example, at least three or four blanks were run at the start of any analysis to ensure system equilibration and before any standard injections. In addition, two blank injections were always done after injection of the highest concentration standard to ensure there was no carry-over from the previous injection. After data acquisition MultiQuant 3.0 software (Sciex, Toronto Canada) was used for linear regression, calculation of r^2^, and plotting of the standard curves. The calibration curve thus obtained allowed for determination of concentration based on peak area directly.

### Calibration curves with internal (heavy peptide) standards

The surrogate peptides having labeled lysine (^13^C_6_ ^15^N_2_) or labeled arginine (^13^C_6_ ^15^N_4_) were synthesized by GenScript to a purity of more than 98% and supplied in a lyophilized form. The lyophilized powder of the peptides was subsequently weighed and reconstituted in LC-MS water having 0.1 % trifluoroacetic acid to a final concentration of 1 mg/mL. A stock cocktail internal standard solution of the labeled peptides was prepared (2 mL) containing a concentration of 2500 ng/mL of each surrogate peptide P, L, N, GP1, GP2 and 12500 ng/mL of surrogate peptide M and GPC in LC-MS grade water having 0.1 % trifluoroacetic acid. For standard curve measurements, a precise (20 µL) amount of the internal standard (labeled) heavy peptide stock was added to each of the dilutions of the light standards. Each external standard solution after addition of the internal standard had a final volume of 1 mL. This resulted in a final concentration of the labeled P, L, N, GP1 and GP2 peptides to be 50 ng/mL whereas the final concentration of the labeled M and GPC peptide was 250 ng/mL. After data acquisition MultiQuant 3.0 software was used for linear regression, calculation of r^2^, and plotting of the standard curves. Since an internal standard was used, the calibration curve thus obtained allowed for determination of concentration based on a peak area ratio.

### Absolute protein concentrations in viral samples with stable isotope labeled internal standard

The heavy labeled internal standard cocktail solution described above was spiked (2.45 µL) into the trypsin treated viral sample (120 µL) to give a final concentration of 50 ng/mL for P, L, N, GP1 and GP2 and 250 ng/mL for M surrogate peptides to match the internal standard concentrations in the external standard solutions. Two samples were taken for each viral batch and each sample was measured in triplicate. The calibration curve obtained from external standards containing internal standards was used to determine absolute concentration of each viral protein in the samples using the MultiQuant 3.0 software.

### Data Analysis

Data analysis was performed using MultiQuant 3.0 (Sciex, Toronto, Canada). The standard curves were generated using linear regression with a weighting of 1/x. A gaussian smooth width of 5.0 points was applied to the peaks at the specific retention times for each of the peptides. The absolute concentrations of the peptides were calculated by the software in ng/mL using a dilution factor of 6. The concentration from MultiQuant in ng/mL was then converted to µM (which corresponded to the full-length viral protein concentrations) using the calculated molecular weight of the surrogate peptides.

### Determination of the Lower Limit of Quantification (LLOQ)

The LLOQ determination was performed according to the ICH (International council for Harmonization) standards. The lowest concentration standard was injected 6 times. LLOQ was then calculated as LLOQ = 10*SD/S, where SD is the standard deviation of the six injections and S is the slope of the standard curve generated for the peptide (45).

### Liquid Chromatography

Liquid chromatography was performed on a NanoAcquity UPLC (Waters, Milford MA) with the column compartment held at 60 °C and maintaining a constant flow rate of 85 µL/min through a C18 reverse phased column (1 mm x 50 mm CSH, 130 Å, 1.7 µ, Waters, Milford MA). Mobile phases A and B were water with 0.1 % formic acid and acetonitrile with 0.1 % formic acid, respectively. In a 45 min run the peptides were separated with a linear gradient of 1-35 % B within the first 35 min, followed by an increase to 90 % B within 5 mins and a re-equilibration.

### Time point cell-infection samples with VSV-GP

The differences in GPC levels were assessed by collecting VSV-GP infected HEK293-F cells at pre-determined time points. Briefly, 5 x 10^5^ HEK293-F cells were plated in 12-well tissue culture plates in BalanCD growth medium (1 mL). Virus stock was diluted to 5 x 10^7^ TCID_50_/mL and infection was carried out at MOI of 1. Mock wells, without viral infection, were included as controls. The cell plates were then incubated in a Kuhner LT-X shaking incubator set to 120 rpm at 37 °C until the intended time of collection. At pre-determined time points (5 min, 3 h and 30 h) the contents of each well were collected into 1.5 mL collection tubes and centrifuged at 300xg for 5 min to pellet the cells. The supernatant was separated without disturbing the cell pellets. The cell pellets were frozen at −80 °C until further processing. Supernatants were collected at the same time points as the cells and were analyzed by tissue culture infectious dose 50% (TCID_50_) assay as reported previously(58).

### Processing of the time point samples

The frozen cell pellets were thawed at room temperature and resuspended in 0.1 % RapiGest diluted in 50 mM ammonium bicarbonate (pH 7.8) containing 1 % sodium deoxycholate (61). The suspension was sonicated for 40 min and then heated to 90 °C for 1 h with 300 rpm shaking before a further 45 min sonication. This was followed by a 10 min centrifugation at 13000 rpm to pellet the cell debris. The supernatant was removed carefully into clean 1.5 mL microcentrifuge tubes. The samples (supernatants) were then reduced by the addition of tris (2-carboxyethyl) phosphine (TCEP) to a final concentration of 20 mM and incubated at 60 °C with 600 rpm shaking for 1 h. Subsequently, they were then cooled to room temperature and alkylated by the addition of iodoacetamide to a final concentration of 40 mM and incubated at 37 °C with 600 rpm shaking for 30 min in the dark. A mixture of trypsin and Endo-Lys-C (Trypsin/Lys-C Mix, Mass Spec Grade, Promega) was used for digestion of the sample. Trypsin/LysC was reconstituted in 50 mM ammonium bicarbonate to a concentration of 1 µg/ml. Samples were digested by addition of 4 µL trypsin, followed by incubation at 37 °C overnight. The digestion was terminated by acidification (addition of trifluoroacetic acid to a final concentration of 0.5% v/v and incubation at 37 °C for 1 h with 600 rpm shaking) followed by centrifugation (13,000xg for 10 min). Prepared samples not run immediately by UPLC-MRM were stored at −20 °C.

### Quantification of the VSV-GP protein levels in the time point samples

For quantification of VSV-GP proteins in the time point samples, the cocktail of the light standards was modified as follows: GP1(60 ng/µL), GP2(80 ng/ µL), GPC (60 ng/ µ L), P (10 ng/ µL), L (10 ng/ µL), N (280 ng/ µL) and M (500 ng/ µL). Serial dilutions of this cocktail stock solution were then prepared to cover the expected concentration range of the peptides. For GP1 and GPC peptides, this dilution range corresponded to 1.5– 1200 ng/mL, 2-1600 ng/mL for GPC, 0.5-200 ng/mL for P, 0.5-100 ng/mL for L, 7.1-5680 ng/mL for N and 25-5000 ng/mL for M. The standard solutions were prepared in approximately 2-fold increments covering the entire concentration range. A stock cocktail internal standard solution of the labeled peptides was prepared (2 mL) containing a concentration of 2500 ng/mL of each surrogate peptide P, L, N, GP1, GP2 and 12500 ng/mL of surrogate peptide M and GPC in LC-MS grade water having 0.1 % trifluoroacetic acid. For standard curve measurements, a precise (20 µL) amount of the internal standard (labeled) heavy peptide stock was added to each of the dilutions of the light standards and to the samples being investigated. Each external standard solution after addition of the internal standard had a final volume of 1 mL. This resulted in a final concentration of the labeled P, L, N, GP1 and GP2 peptides to be 50 ng/mL whereas the final concentration of the labeled M and GPC peptide was 250 ng/mL.

### Mass Spectrometry

All MRM experiments were carried out on a triple stage quadrupole mass spectrometer (6500 QTrap^®^, Sciex) using a Turbo Spray Ion Drive source in positive ion mode. Analyst Software (version 7.1) was used for data acquisition with ion spray voltage set at 5200 V, curtain gas 30, temperature 300 ⁰C, and both Gas 1 and 2 to 30 psi. Collision gas was set to 11 psi. Declustering, entrance, and cell exit potentials were 80, 10, and 18 V, respectively (**Table S1**). Both quadrupoles Q1 and Q3 were set to unit resolution. The dwell time for each transition was adjusted to 10 ms to maintain a cycle time at or near 0.3 s to provide good delineation of the UPLC peaks. Scheduled MRM was not used since all surrogate peptide peaks were well resolved by the UPLC gradient method.

### Stability Analysis

The standard stock solutions of the peptides were stored at 4 °C and run after a span of 7 days to generate calibration curves as described above. The assessment of stability of the stock solutions were done by analysis of the r^2^ value and the slope of the curve. The peptide stock was considered undergoing degradation over time if, r^2^ value was less than 0.98 or the slope changed when compared to freshly prepared stock solutions. Long term stability was evaluated by storage of the standard stock solutions at −25 °C and analyzing the standard curves over a span of 3 weeks.

## Supporting information

Supplementary Information Absolute Quantification of Viral Proteins in pseudo typed Vesicular Stomatitis Virus (VSV-GP) using Ultra High-Performance L

## Abbreviations

LCMV: Lymphocytic Choriomeningitis Virus
LLOQ: Lower Limit of Quantification
MRM: Multiple Reaction monitoring
qRT: PCR-Quantitative reverse transcriptase polymerase chain reaction
TCID50: 50% Tissue Culture Infectious Dose
UPLC: Ultra high-performance liquid chromatography
VSV: Vesicular Stomatitis Virus
VSV: GP-Pseudotyped VSV with LCMV envelope protein.

## ASSOCIATED CONTENT

### Supporting Information

The supporting information is available.

Supplementary figures for chromatograms, standard curves and time point protein samples are attached in word document.

Supplementary information for all transitions of surrogate peptides is attached in the excel sheet.

### Notes

The authors declare no competing financial interests. All authors were employed by Boehringer Ingelheim Pharmaceuticals Inc. (BIPI) at the time of the experiments.

## REFERENCES

1. Sanyal G, Särnefält A, Kumar A. 2021. Considerations for bioanalytical characterization and batch release of COVID-19 vaccines. Npj Vaccines 6:53.

2. González-Domínguez I, Puente-Massaguer E, Cervera L, Gòdia F. 2020. Quality Assessment of Virus-Like Particles at Single Particle Level: A Comparative Study. Viruses 12:223.

3. Lothert K, Eilts F, Wolff MW. 2022. Quantification methods for viruses and virus-like particles applied in biopharmaceutical production processes. Expert Rev Vaccines 21:1029– 1044.

4. Lupberger J, Croonenborghs T, Suarez AAR, Renne NV, Jühling F, Oudot MA, Virzì A, Bandiera S, Jamey C, Meszaros G, Brumaru D, Mukherji A, Durand SC, Heydmann L, Verrier ER, Saghire HE, Hamdane N, Bartenschlager R, Fereshetian S, Ramberger E, Sinha R, Nabian M, Everaert C, Jovanovic M, Mertins P, Carr SA, Chayama K, Dali-Youcef N, Ricci R, Bardeesy NM, Fujiwara N, Gevaert O, Zeisel MB, Hoshida Y, Pochet N, Baumert TF. 2019. Combined Analysis of Metabolomes, Proteomes, and Transcriptomes of Hepatitis C Virus– Infected Cells and Liver to Identify Pathways Associated With Disease Development. Gastroenterology 157:537–551.e9.

5. Gao Y, Fillmore TL, Munoz N, Bentley GJ, Johnson CW, Kim J, Meadows JA, Zucker JD, Burnet MC, Lipton AK, Bilbao A, Orton DJ, Kim Y-M, Moore RJ, Robinson EW, Baker SE, Webb-Robertson B-JM, Guss AM, Gladden JM, Beckham GT, Magnuson JK, Burnum-Johnson KE. 2020. High-Throughput Large-Scale Targeted Proteomics Assays for Quantifying Pathway Proteins in Pseudomonas putida KT2440. Frontiers Bioeng Biotechnology 8:603488.

6. Picotti P, Aebersold R. 2012. Selected reaction monitoring–based proteomics: workflows, potential, pitfalls and future directions. Nat Methods 9:555–566.

7. Kondrat RW, Cooks RG. 1978. Direct Analysis of Mixtures by Mass Spectrometry. Analytical Chemistry 50:81A–92A.

8. Yost RA, Enke CG. 1979. Triple quadrupole mass spectrometry for direct mixture analysis and structure elucidation. Anal Chem 51:1251–1264.

9. Sherwood CA, Eastham A, Lee LW, Risler J, Mirzaei H, Falkner JA, Martin DB. 2009. Rapid Optimization of MRM-MS Instrument Parameters by Subtle Alteration of Precursor and Product m/z Targets. J Proteome Res 8:3746–3751.

10. Janecki DJ, Bemis KG, Tegeler TJ, Sanghani PC, Zhai L, Hurley TD, Bosron WF, Wang M. 2007. A multiple reaction monitoring method for absolute quantification of the human liver alcohol dehydrogenase ADH1C1 isoenzyme. Anal Biochem 369:18–26.

11. Matraszek-Zuchowska I, Wozniak B, Posyniak A. 2016. Comparison of the Multiple Reaction Monitoring and Enhanced Product Ion Scan Modes for Confirmation of Stilbenes in Bovine Urine Samples Using LC–MS/MS QTRAP® System. Chromatographia 79:1003–1012.

12. Ulintz PJ, Yocum AK, Bodenmiller B, Aebersold R, Andrews PC, Nesvizhskii AI. 2009. Comparison of MS2-Only, MSA, and MS2/MS3 Methodologies for Phosphopeptide Identification. J Proteome Res 8:887–899.

13. A K Michael, Derek S, Juncong Y, J C Tyra, M J Angela, B H Darryl, Leigh A N, H B Christoph. Multiple reaction monitoring-based, multiplexed, absolute quantitation of 45 proteins in human plasma. Mol Cell Proteomics 8:1860–1877.

14. Peterson AC, Russell JD, Bailey DJ, Westphall MS, Coon JJ. 2012. Parallel Reaction Monitoring for High Resolution and High Mass Accuracy Quantitative, Targeted Proteomics*. Mol Cell Proteomics 11:1475–1488.

15. Kirkpatrick DS, Gerber SA, Gygi SP. 2005. The absolute quantification strategy: a general procedure for the quantification of proteins and post-translational modifications. Methods 35:265–273.

16. Mayya V, Rezual K, Wu L, Fong MB, Han DK. 2006. Absolute Quantification of Multisite Phosphorylation by Selective Reaction Monitoring Mass Spectrometry Determination of Inhibitory Phosphorylation Status of Cyclin-Dependent Kinases* S. Mol Cell Proteomics 5:1146–1157.

17. Janecki DJ, Bemis KG, Tegeler TJ, Sanghani PC, Zhai L, Hurley TD, Bosron WF, Wang M. 2007. A multiple reaction monitoring method for absolute quantification of the human liver alcohol dehydrogenase ADH1C1 isoenzyme. Anal Biochem 369:18–26.

18. Gerber SA, Kettenbach AN, Rush J, Gygi SP. 2007. Quantitative Proteomics by Mass Spectrometry. Methods Mol Biology 359:71–86.

19. Gerber SA, Rush J, Stemman O, Kirschner MW, Gygi SP. 2003. Absolute quantification of proteins and phosphoproteins from cell lysates by tandem MS. Proc National Acad Sci 100:6940–6945.

20. Salvatore S, Gerber SA, Kettenbach AN, Rush J, Gygi SP. 2006. Quantitative Proteomics by Mass Spectrometry 71–86.

21. Riepler L, Frommelt L-S, Wilmschen-Tober S, Mbuya W, Held K, Volland A, Laer D von, Geldmacher C, Kimpel J. 2023. Therapeutic Efficacy of a VSV-GP-based Human Papilloma Virus Vaccine in a Murine Cancer Model. J Mol Biol 435:168096.

22. Tisoncik-Go J, Voss KM, Lewis TB, Muruato AE, Kuller L, Finn EE, Betancourt D, Wangari S, Ahrens J, Iwayama N, Grant RF, Murnane RD, Edlefsen PT, Fuller DH, Barber GN, Gale M, O’Connor MA. 2023. Evaluation of the immunogenicity and efficacy of an rVSV vaccine against Zika virus infection in macaca nemestrina. Front Virol 3:1108420.

23. Muik A, Stubbert LJ, Jahedi RZ, Geiβ Y, Kimpel J, Dold C, Tober R, Volk A, Klein S, Dietrich U, Yadollahi B, Falls T, Miletic H, Stojdl D, Bell JC, Laer D von. 2014. Re-engineering Vesicular Stomatitis Virus to Abrogate Neurotoxicity, Circumvent Humoral Immunity, and Enhance Oncolytic Potency. Cancer Res 74:3567–3578.

24. Tober R, Banki Z, Egerer L, Muik A, Behmüller S, Kreppel F, Greczmiel U, Oxenius A, Laer D von, Kimpel J. 2014. VSV-GP: a Potent Viral Vaccine Vector That Boosts the Immune Response upon Repeated Applications. J Virol 88:4897–4907.

25. Abraham G, Banerjee AK. 1976. Sequential transcription of the genes of vesicular stomatitis virus. Proc National Acad Sci 73:1504–1508.

26. Raux H, Obiang L, Richard N, Harper F, Blondel D, Gaudin Y. 2010. The Matrix Protein of Vesicular Stomatitis Virus Binds Dynamin for Efficient Viral Assembly. J Virol 84:12609– 12618.

27. Wagner RR, Schnaitman TA, Snyder RM. 1969. Structural Proteins of Vesicular Stomatitis Viruses. J Virol 3:395–403.

28. Green TJ, Luo M. 2009. Structure of the vesicular stomatitis virus nucleocapsid in complex with the nucleocapsid-binding domain of the small polymerase cofactor, P. Proc National Acad Sci 106:11713–11718.

29. Jenni S, Bloyet L-M, Diaz-Avalos R, Liang B, Whelan SPJ, Grigorieff N, Harrison SC. 2020. Structure of the Vesicular Stomatitis Virus L Protein in Complex with Its Phosphoprotein Cofactor. Cell Reports 30:53–60.e5.

30. Wagner RR, Schnaitman TC, Snyder RM, Schnaitman CA. 1969. Protein Composition of the Structural Components of Vesicular Stomatitis Virus. J Virol 3:611–618.

31. Bosma B, Plessis F du, Ehlert E, Nijmeijer B, Haan M de, Petry H, Lubelski J. 2018. Optimization of viral protein ratios for production of rAAV serotype 5 in the baculovirus system. Gene Ther 25:415–424.

32. Bederka LH, Bonhomme CJ, Ling EL, Buchmeier MJ. 2014. Arenavirus Stable Signal Peptide Is the Keystone Subunit for Glycoprotein Complex Organization. Mbio 5:e02063–14.

33. Saunders AA, Ting JPC, Meisner J, Neuman BW, Perez M, Torre JC de la, Buchmeier MJ. 2007. Mapping the Landscape of the Lymphocytic Choriomeningitis Virus Stable Signal Peptide Reveals Novel Functional Domains. J Virol 81:5649–5657.

34. Hastie KM, Igonet S, Sullivan BM, Legrand P, Zandonatti MA, Robinson JE, Garry RF, Rey FA, Oldstone MB, Saphire EO. 2016. Crystal structure of the prefusion surface glycoprotein of the prototypic arenavirus LCMV. Nat Struct Mol Biol 23:513–521.

35. Pennington HN, Lee J. 2022. Lassa virus glycoprotein complex review: insights into its unique fusion machinery. Bioscience Rep 42:BSR20211930.

36. Kunz S, Edelmann KH, Torre J-C de la, Gorney R, Oldstone MBA. 2003. Mechanisms for lymphocytic choriomeningitis virus glycoprotein cleavage, transport, and incorporation into virions. Virology 314:168–178.

37. Adams KJ, Pratt B, Bose N, Dubois LG, John-Williams LS, Perrott KM, Ky K, Kapahi P, Sharma V, MacCoss MJ, Moseley MA, Colton CA, MacLean BX, Schilling B, Thompson JW, Consortium ADM. 2020. Skyline for Small Molecules: A Unifying Software Package for Quantitative Metabolomics. J Proteome Res 19:1447–1458.

38. Lange V, Picotti P, Domon B, Aebersold R. 2008. Selected reaction monitoring for quantitative proteomics: a tutorial. Mol Syst Biol 4:222–222.

39. Gerber SA, Rush J, Stemman O, Kirschner MW, Gygi SP. 2003. Absolute quantification of proteins and phosphoproteins from cell lysates by tandem MS. Proc National Acad Sci 100:6940–6945.

40. Stahl-Zeng J, Lange V, Ossola R, Eckhardt K, Krek W, Aebersold R, Domon B. 2007. High Sensitivity Detection of Plasma Proteins by Multiple Reaction Monitoring of N-Glycosites*. Mol Cell Proteomics 6:1809–1817.

41. Wang C, Wang Y, Xie H, Zhan C, He X, Liu R, Hu R, Shen J, Jia Y. 2022. Establishment and validation of an SIL-IS LC–MS/MS method for the determination of ibuprofen in human plasma and its pharmacokinetic study. Biomed Chromatogr 36:e5287.

42. Thomas D, Newcomb WW, Brown JC, Wall JS, Hainfeld JF, Trus BL, Steven AC. 1985. Mass and molecular composition of vesicular stomatitis virus: a scanning transmission electron microscopy analysis. J Virol 54:598–607.

43. Barge A, Gaudin Y, Coulon P, Ruigrok RW. 1993. Vesicular stomatitis virus M protein may be inside the ribonucleocapsid coil. J Virol 67:7246–7253.

44. Soh TK, Whelan SPJ. 2015. Tracking the Fate of Genetically Distinct Vesicular Stomatitis Virus Matrix Proteins Highlights the Role for Late Domains in Assembly. J Virol 89:11750– 11760.

45. ICH guideline M10 on bioanalytical method validation and study sample analysis.

46. Zhang H, Liu Q, Zimmerman LJ, Ham A-JL, Slebos RJC, Rahman J, Kikuchi T, Massion PP, Carbone DP, Billheimer D, Liebler DC. 2011. Methods for Peptide and Protein Quantitation by Liquid Chromatography-Multiple Reaction Monitoring Mass Spectrometry*. Mol Cell Proteomics 10:M110.006593.

47. Ball LA, Pringle CR, Flanagan B, Perepelitsa VP, Wertz GW. 1999. Phenotypic Consequences of Rearranging the P, M, and G Genes of Vesicular Stomatitis Virus. J Virol 73:4705–4712.

48. Zhong X, Nayak S, Guo L, Raidas S, Zhao Y, Weiss R, Andisik M, Elango C, Sumner G, Irvin SC, Partridge MA, Yan H, E SY, Qiu H, Mao Y, Torri A, Li N. 2021. Liquid Chromatography-Multiple Reaction Monitoring-Mass Spectrometry Assay for Quantitative Measurement of Therapeutic Antibody Cocktail REGEN-COV Concentrations in COVID-19 Patient Serum. Anal Chem 93:acs.analchem.1c01613.

49. Lenz O, Meulen J ter, Klenk H-D, Seidah NG, Garten W. 2001. The Lassa virus glycoprotein precursor GP-C is proteolytically processed by subtilase SKI-1/S1P. Proc National Acad Sci 98:12701–12705.

50. Branco LM, Garry RF. 2009. Characterization of the Lassa virus GP1 ectodomain shedding: implications for improved diagnostic platforms. Virol J 6:147.

51. Schlie K, Maisa A, Lennartz F, Ströher U, Garten W, Strecker T. 2010. Characterization of Lassa Virus Glycoprotein Oligomerization and Influence of Cholesterol on Virus Replication. J Virol 84:983–992.

52. Illick MM, Branco LM, Fair JN, Illick KA, Matschiner A, Schoepp R, Garry RF, Guttieri MC. 2008. Uncoupling GP1 and GP2 expression in the Lassa virus glycoprotein complex: implications for GP1 ectodomain shedding. Virol J 5:161.

53. Liigand P, Kaupmees K, Kruve A. 2019. Influence of the amino acid composition on the ionization efficiencies of small peptides. Journal of Mass Spectrometry 54:481–487.

54. Redondo N, Madan V, Alvarez E, Carrasco L. 2015. Impact of Vesicular Stomatitis Virus M Proteins on Different Cellular Functions. Plos One 10:e0131137.

55. Rosen CA, Cohen PS, Ennist HL. 1983. Identification of a new protein present in vesicular stomatitis virus-infected Chinese hamster ovary cells as a degradation product of viral M protein. Virology 130:331–341.

56. Liang B, Li Z, Jenni S, Rahmeh AA, Morin BM, Grant T, Grigorieff N, Harrison SC, Whelan SPJ. 2015. Structure of the L Protein of Vesicular Stomatitis Virus from Electron Cryomicroscopy. Cell 162:314–327.

57. Jenni S, Horwitz JA, Bloyet L-M, Whelan SPJ, Harrison SC. 2022. Visualizing molecular interactions that determine assembly of a bullet-shaped vesicular stomatitis virus particle. Nat Commun 13:4802.

58. Dambra R, Matter A, Graca K, Akhand SS, Mehta S, Bell-Cohn A, Swenson JM, Abid S, Xin D, Lewis C, Coyle L, Wang M, Bunosso K, Maugiri M, Ruiz R, Cirillo CM, Fogal B, Grimaldi C, Vigil A, Wood C, Ashour J. 2023. Nonclinical pharmacokinetics and biodistribution of VSV-GP using methods to decouple input drug disposition and viral replication. Mol Ther - Methods Clin Dev 28:190–207.

59. Gautam S, Xin D, Garcia AP, Spiesschaert B. 2022. Single-step rapid chromatographic purification and characterization of clinical stage oncolytic VSV-GP. Frontiers Bioeng Biotechnology 10:992069.

60. Silva JC, Gorenstein MV, Li G-Z, Vissers JPC, Geromanos SJ. 2006. Absolute Quantification of Proteins by LCMSE A Virtue of Parallel ms Acquisition * S. Mol Cell Proteomics 5:144–156.

61. Lin Y, Zhou J, Bi D, Chen P, Wang X, Liang S. 2008. Sodium-deoxycholate-assisted tryptic digestion and identification of proteolytically resistant proteins. Anal Biochem 377:259–266.

