## Supplementary Information Absolute Quantification of Viral Proteins in pseudo typed Vesicular Stomatitis Virus (VSV-GP) using Ultra High-Performance L for "Absolute Quantification of Viral Proteins from Pseudotyped Vesicular Stomatitis Virus (VSV-GP) using Ultra High-Performance Liquid Chromatography- Multiple Reaction Monitoring (UPLC-MRM)"

♣ Materials and Analytical Sciences, * Drug Metabolism and Pharmacokinetics, Boehringer Ingelheim Pharmaceuticals, 900 Ridgebury Road, Ridgefield, Connecticut – 06877

♦ Therapeutic Virus Development Group, Virus Therapeutic Center, Boehringer Ingelheim Pharma GmbH & Co. KG, Birkendorfer Str. 65, 88400 Biberach

-

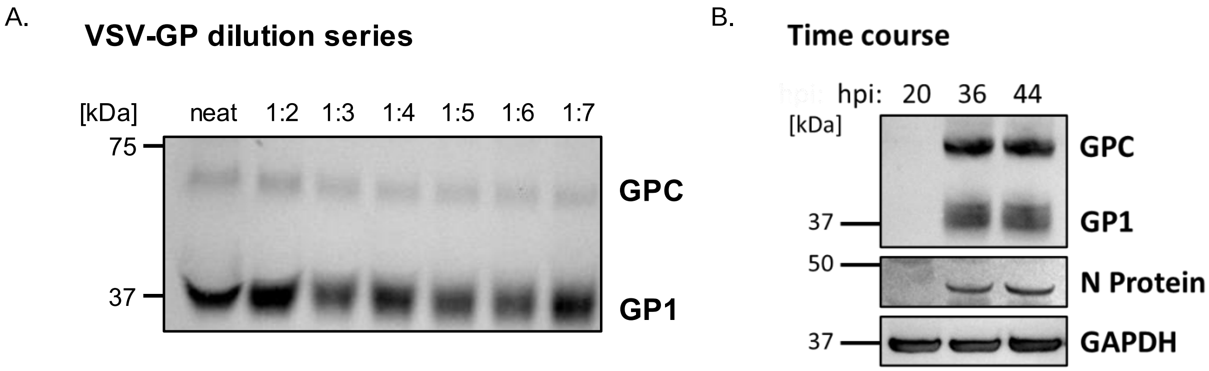

**Fig. S1: GP1 detection with SDS-PAGE / western blot.** To assess the maturation state of the glycoprotein either drug product or lysate samples were analyzed on a western blot with a GP1 directed antibody. A) A dilution series with a VSV-GP sample after downstream processing was performed. Two bands at the corresponding molecular weights for GP1 and GPC were detected for pure virus and the employed dilutions. B) To determine the intracellular ratio of GPC to GP1 in lysates of HEK293F, cells were infected with VSV-GP at an MOI of 0.0005 and harvested at different timepoints after infection. While the amount of virus after 20 hours post infection (hpi) was probably too low to detect GP1 or N with western blot two bands corresponding to the molecular weight of GPC and GP1 can be seen at 36 hpi and 44 hpi.

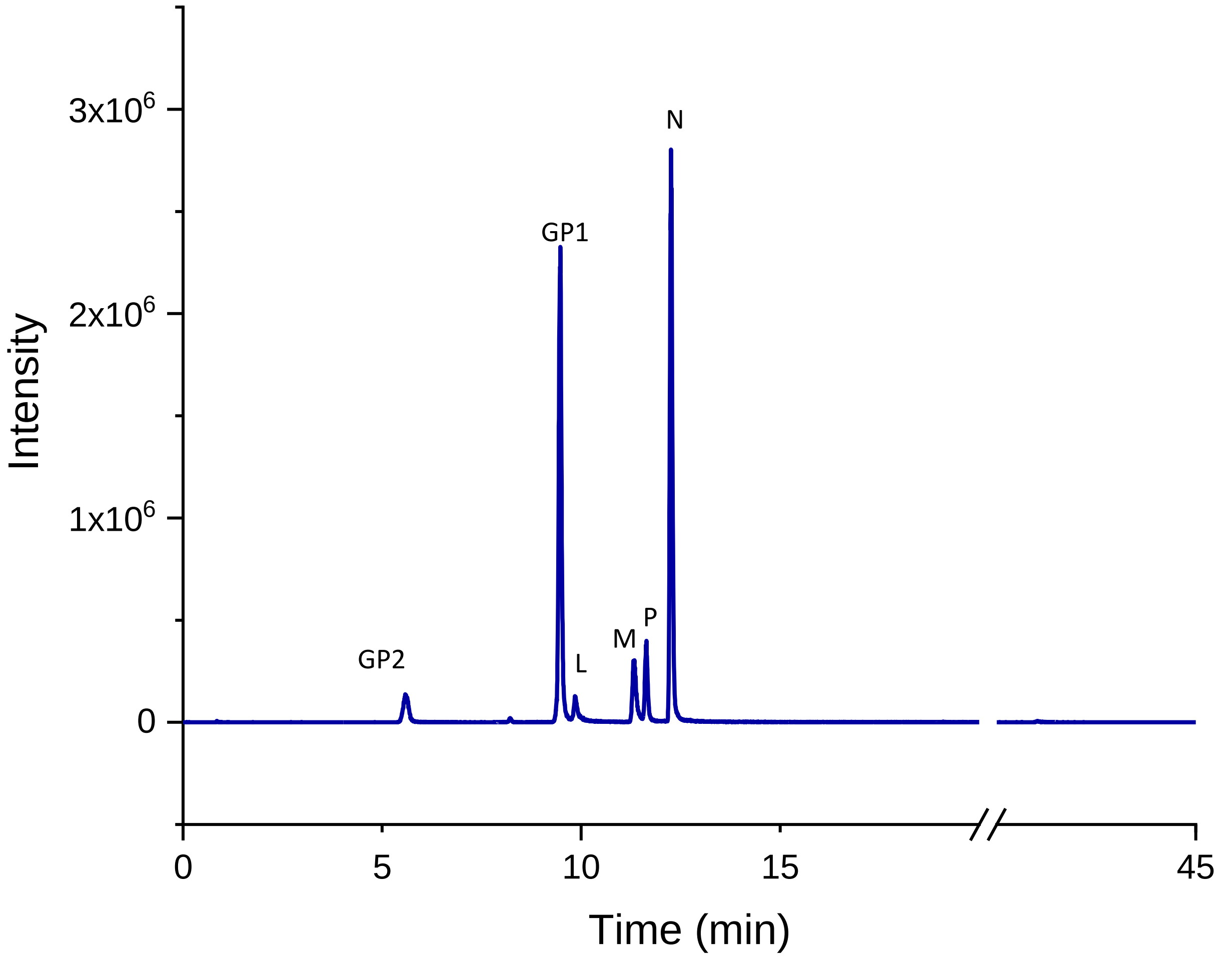

**Fig. S2: Total ion chromatogram (TIC) showing peaks of all the VSV-GP proteins detected:** A Waters NanoAcquity UPLC was used to separate the peaks in a 45 min run, with retention times retention time 5.71 min (GP2), 9.49 min (GP1) corresponds to GP1, 9.85 min (L), 11.32 min (M), 11.71 min (P) and 12.26 min (N). The values of the retention time and corresponding transitions for the peptides have been reported in the main text. The TIC has been extracted from one of the standard curve dilution samples having 250 ng/mL GP1 and GP2, 25 ng/mL P and L, 710 ng/mL N and 1250 ng/mL M light unlabeled peptide concentration, 50 ng/mL of GP1, GP2, P, L, N and 250 ng/mL heavy labeled peptide concentrations. The TIC was extracted with Peak View 3.0 and graphs were replotted with Origin 2019.

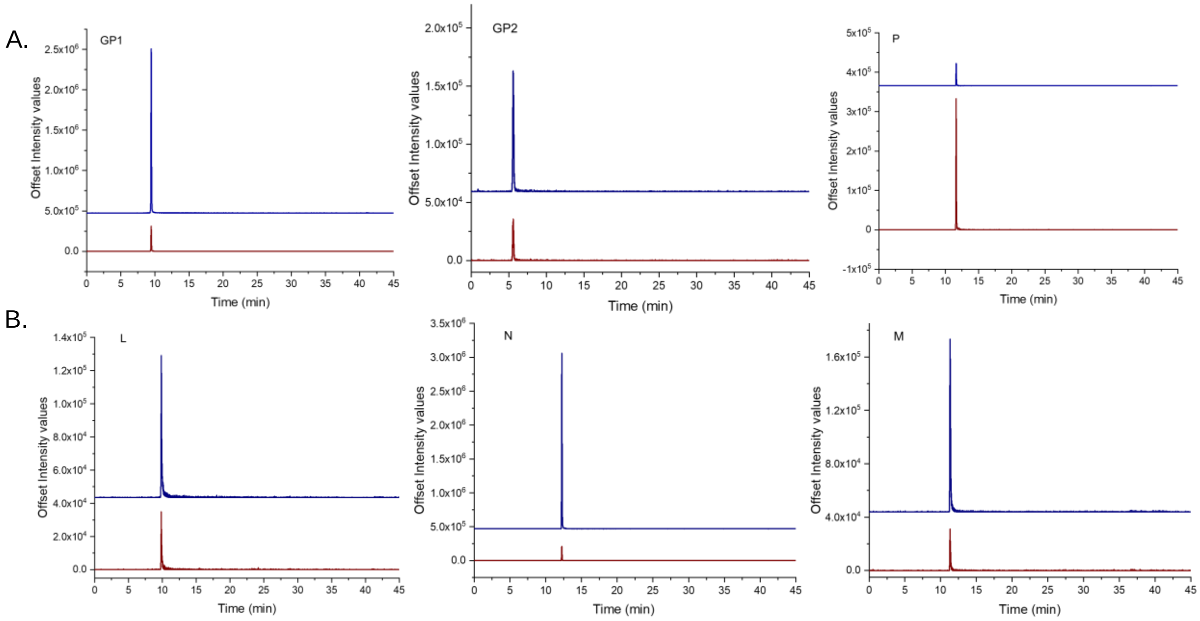

**Fig. S3: Extracted ion chromatogram (XIC) of each surrogate peptide (both unlabeled and labeled) corresponding to VSV-GP proteins.** A). XIC of unlabeled and labeled surrogate peptides corresponding to GP1, GP2 and P. The labeled surrogates for these proteins had a heavy label at N-terminal lysine residue ($^{13}$C6, $^{15}$N2) resulting in a mass shift of 8 Da or N-terminal arginine label ($^{13}$C6, $^{15}$N4) with a mass shift of 10 Da. They elute at the same retention times of 9.49 min (GP1), 5.71 min (GP2) and 11.71 min (P) providing specificity of identification for the peptides. The blue trace corresponds to the unlabeled surrogate peptide while the red trace corresponds to the heavy labeled surrogate peptide. B). XIC of unlabeled and labeled surrogate peptides corresponding to L, N and M. The labeled surrogates for these proteins had a heavy label at N-terminal lysine residue ($^{13}$C6, $^{15}$N2) for N and M resulting in mass shift of 8 Da, heavy labeled at N-terminal arginine residue ($^{13}$C6,$^{15}$N4) for surrogate peptide L resulting in a mass shift of 10 Da. They elute at the same retention times of 9.85 min (L), 12.26 min (N) and 11.32 min (M)

providing specificity of identification for the peptides. The blue trace corresponds to the unlabeled surrogate peptide while the red trace corresponds to the heavy labeled surrogate peptide. Each of the XIC peaks have been extracted from one of the standard curve dilution samples having 250 ng/mL GP1 and GP2, 25 ng/mL P and L, 710 ng/mL N and 1250 ng/mL M light unlabeled peptide concentration, 50 ng/mL of GP1, GP2, P, L, N and 250 ng/mL heavy labeled peptide concentrations. The XIC was extracted with Peak View 3.0 and graphs were replotted with Origin 2019.

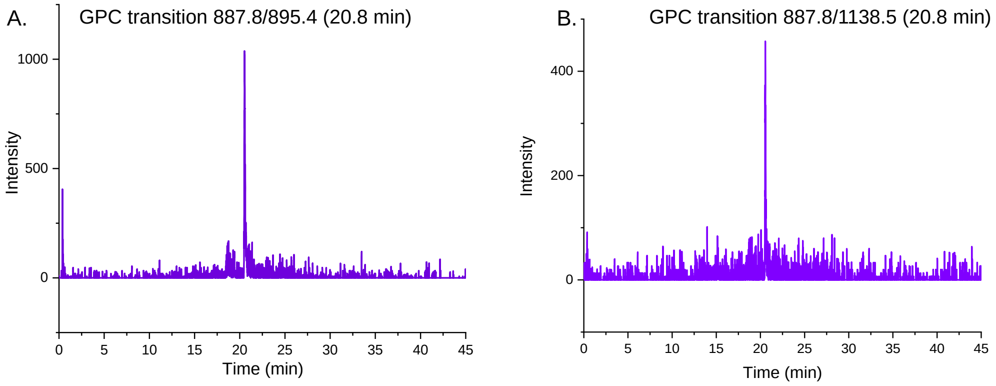

C.

MGQIVTMFEALPHIIDEVINIVIIVLIIITSIKAVYNFATCGILALVSFLFLAGRSCG**MY**
**GLNGPDIYKGVYQFKSVEFDMSHLNLTMPNACSANNSHHYISMGSSGLELTFTNDSILNH**
**NFCNLTSAFNKKTFDHTLMSIVSSLHLSIRGNSNHKAVSCDFNNGITIQYNLSFSDPQSA**
**ISQCRTFRGRVLDMFRTAFGGKYMRSGWGWAGSDGKTTWCSQTSYQYLIIQNRTWENHCR**
**YAGPFGMSRILFAQEKTKFLTRR*LAGTFTWTLSDSSGVENPGGYCLTK*WMILAAELKCFG**
**NTAVAKCNVNHDEEFCDMLRLIDYNKAALSKFKQDVESALHVFKTTVNSLISDQLLMRNH**
**LRDLMGVPYCNYSKFWYLEHAKTGETSVPKCWLVTNGSYLNETHFSDQIEQEADNMITEM**
**LRKDYIKRQGSTPLALMDLLMFSTSAYLISIFLHLVKIPTHRHIKGGSCPKPHRLTNKGI**
**CSCGAFKVPGVKTIWKRR**

**Fig. S4: Transitions of surrogate peptide for full-length GPC:** The identified peptide LAGTFTWTLSDSSGVENPGGYC[CAM]LTK surrogate for full length GPC eluted at retention time 20.8 min and two transitions were identified at the same retention time, providing confidence in the detection of the surrogate peptide specific for GPC. A) Chromatogram at 20.8 min for transition 887.8/895.4. B) Chromatogram for transition 887.8/1138.5. C) Full-length GPC sequence showing surrogate peptide selection: The surrogate peptide selection for GPC containing both GP1 and GP2 (peptide sequence highlighted in yellow). The sequence in red indicates sequence corresponding to GP1, the sequence in green represents sequence corresponding to GP2 (1).

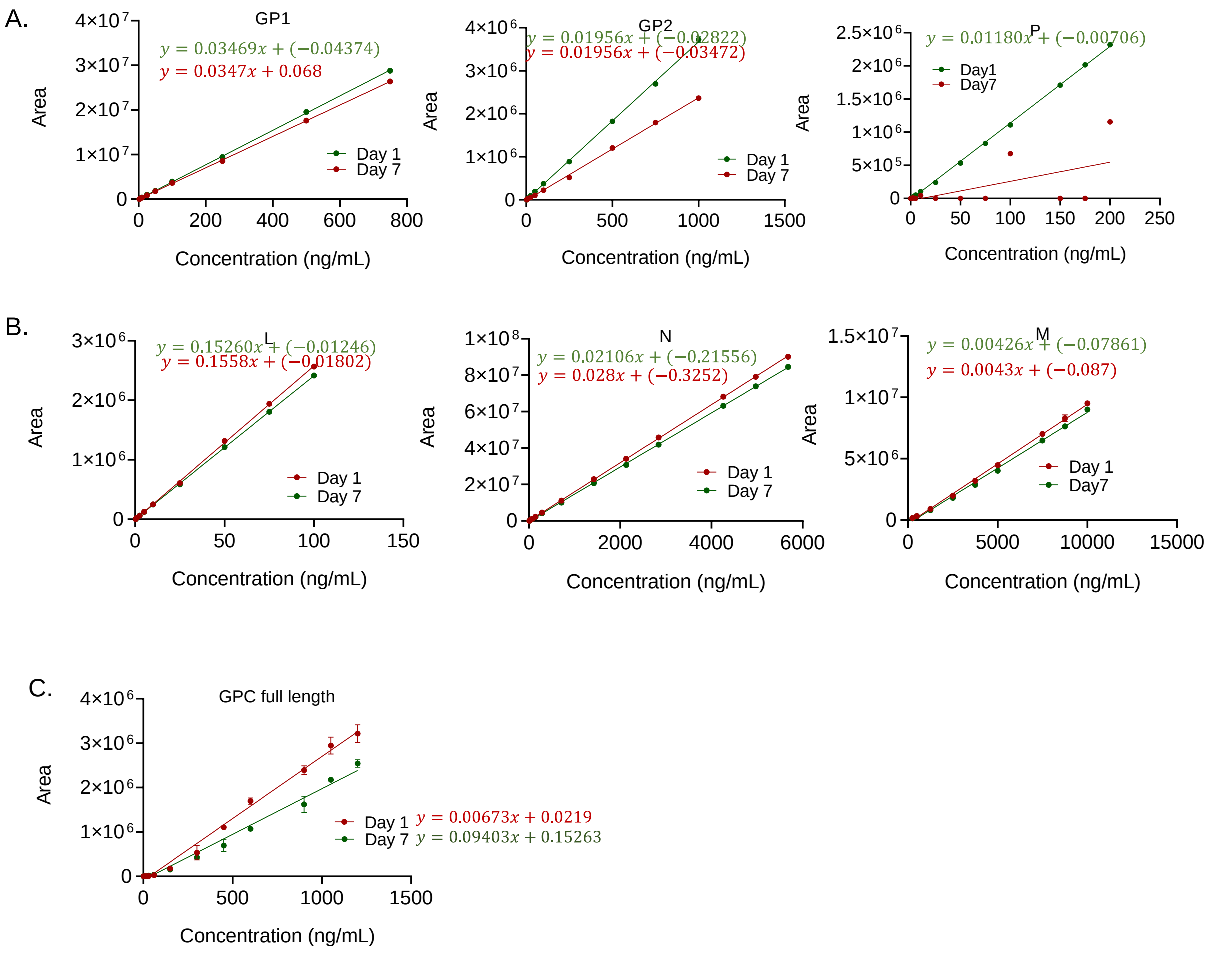

**Fig. S5: Stability study with surrogate peptide standards:** The standard solutions (dilutions) of the peptides were prepared and stored at 4 °C and evaluated for standard curves at the 7th day. The figure shows comparison of the standard curves from freshly prepared standards (red line connecting red dots) vs standard curves from the run done at the 7th day (green line connecting green dots). The equation for linear regression written in red represents one for freshly prepared standards and the equation written in green represents one for standards run at the 7th day A) Shows standard curves for GP1, GP2 and P B) Shows standard curves for L, N and M and C) shows standard curves for GPC respectively. The linear regression analysis of the standard curves has been done in MultiQuant 3.0 using a weighting of 1/x.

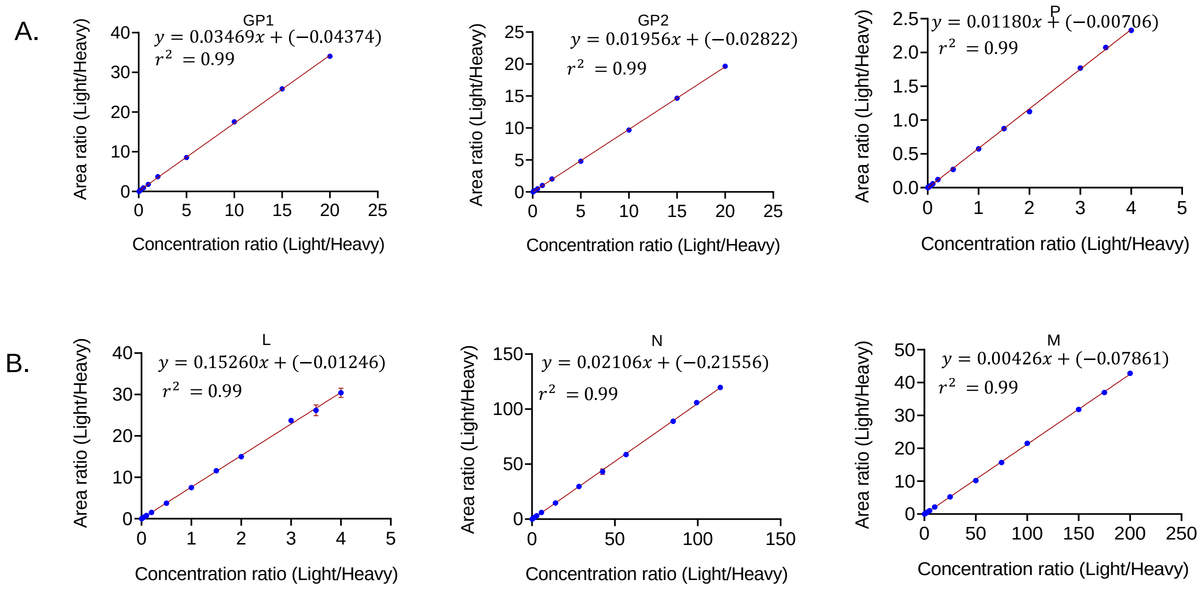

**Fig. S6: Standard curves from unlabeled surrogate peptide standard solutions containing labeled (heavy) internal standards: A, B.** Standard curves from standard solutions containing labeled (heavy) surrogate peptide internal standards corresponding to each protein, labeled at N-terminal lysine residue ($^{13}C_6$,$^{15}N_2$) or N-terminal arginine residue ($^{13}C_6$,$^{15}N_4$). The heavy labeled surrogate peptides were spiked into the cocktail of light standards at a known and fixed concentration which was 50 ng/mL for peptides GP1, GP2, P, L and N and 250 ng/mL for peptide M. The concentration range of the light standards for each of the peptide had the same concentration range as used for unlabeled surrogates (Figure 1). The standard curve was plotted in MultiQuant 3.0 using linear regression and a weighting 1/x as peak area ratios (y-axis) vs concentration ratios of light/heavy (x-axis). **A** shows plots for GP1, GP2 and P, **B** shows plots for L, N and M respectively. Each linear regression (standard curve) was generated from three independent runs with duplicate injections thus creating an average of six data points for each concentration.

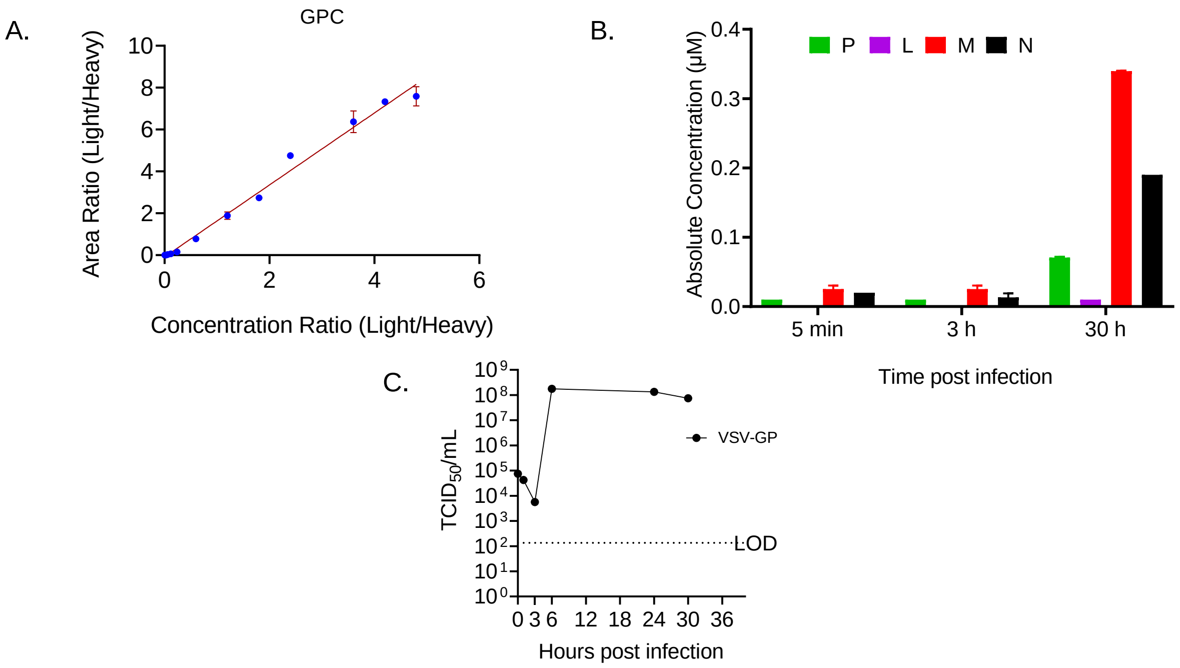

**Fig. S7: Comparison of VSV-GP proteins in time point lysates of VSV-GP HEK-293 infected cells**: A) Standard curve with the GPC heavy surrogate peptide labeled at N-terminal lysine residue ($^{13}C_6$,$^{15}N_2$) at a fixed concentration of 250 ng/mL B) Comparison of absolute concentrations (µM) of VSV-GP proteins (P, L, M and N) in 5 min, 3 h and 30 h time point lysates collected from VSV-GP infected HEK-293F cells. The plotted histograms are the mean of two samples from each time point with each sample run in triplicates, error bars representing standard deviation from mean (**Table 5**). C) Tissue culture infective dose 50% ($TCID_{50}$) was performed on supernatants collected at same time points (5 min, 3 h and 30 h) to quantify infectious virus progeny release from cells confirming the complete viral life cycle.

| Table S1: MRM parameters of surrogate peptides for the VSV-GP proteins in the study | | | | | | |
|---|---|---|---|---|---|---|
| **Viral protein** | **Target peptide[a]** | **Collision energy** | **Declustering potential (eV)** | **Entrance potential (eV)** | **Cell exit potential(eV)** | **Dwell Time (ms)** |
| **GP1** | ILFAQEK | 19.8 | 80 | 10 | 18 | 10 |
| **GP2** | C[CAM]FGNTAVAK | 22.7 | 80 | 10 | 18 | 10 |
| **P** | SQWLSTIK | 22.6 | 80 | 10 | 18 | 10 |
| **L** | EFLNPDER | 22.0 | 80 | 10 | 18 | 10 |
| **N** | IIDNTVVPK | 25.9 | 80 | 10 | 18 | 10 |
| **M** | RFNIGLYK | 34.2 | 80 | 10 | 18 | 10 |
| **GPC** | LAGTFTWTLSDSSGVENPGGYC[CAM]LTK | 40.6 | 80 | 10 | 18 | 10 |

[a] . The parameters tabulated here are same for both light and heavy surrogate peptides.

| Table S2: Genome content and Mean $TCID_{50}$/mL for different purified viral batches | | |
|---|---|---|
| **Viral Batch** | **Genome Content/mL (GC/mL) [a]** | **Mean $TCID_{50}$/ mL [b]** |
| **Batch 1** | $3.90x10^{11}$ | $2.80x10^{10}$ |
| **Batch 2** | $3.19x10^{11}$ | $8.91x1^{09}$ |
| **Batch 3** | $4.01x10^{11}$ | $1.06x10^{10}$ |
| **Batch 4** | $6.94x10^{11}$ | $9.75x10^{9}$ |
| **Batch 5** | $1.02x10^{12}$ | $1.05x10^{10}$ |

| | | |
|---|---|---|
| **Batch 6** | $6.89 \times 10^{11}$ | $6.34 \times 10^{9}$ |
| **Batch 7** | $7.93 \times 10^{11}$ | $7.32 \times 10^{9}$ |
| **Batch 8** | $8.23 \times 10^{11}$ | $9.76 \times 10^{9}$ |

[a]. The genome content was determined by qRT-PCR.

[b]. The mean $TCID_{50}$/mL value provided information on the viral titer which was equivalent to the concentration of the virus.

**Supplementary Methods:**

**Viral Nucleic Acid extraction and qRT-PCR for viral genome quantification**

Viral nucleic acid was manually extracted and purified from virus stocks and then the extracted viral samples were subjected to one-step qRT-PCR using the method previously described by Dambra et al. (2) .

**SDS-PAGE / western blot**

Virus samples were diluted with PBS and 4x Lämmli buffer (supplied with β-mercaptoethanol 1:10) was added to 1x. The mix was vortexed and heated to 95 °C for 5 min with shaking. For cell lysis, 1*10^6 cells were spun down at 1000 g for 5 min. Cells were washed and 50 µL of lysis buffer with protease inhibitor (Cocktail, Roche) was added to the cell pellet followed by 20 min incubation at 4°C. Lysed cells were spun down at 21,300 g for 15 min at 4°C. The supernatants were transferred to a new tube to which 4x Lämmli buffer (supplied with β-mercaptoethanol 1:10) was added. Further, samples were heated to 95°C for 5 min. Proteins were separated by SDS-PAGE (NuPage Bis-Tris 10 %, InVitrogen) and blotted on a nitrocellulose membrane (Trans-Blot Turbo 0,2 µm pore size, BIO-RAD). After transfer, membranes were blocked in 1 % casein blocking buffer for 30 min. Antibodies were diluted to 1 : 2,000 (Anti-GP KL25-3A7, Syngene), 1 : 1,000 (Anit-VSV N [19G4], Absolute Biotech Company) and 1 : 2,000 (GAPDH, Cell Signaling Technology). Following, membranes were incubated over night with the primary antibodies at 4 °C. Membranes were washed and secondary antibodies conjugated to horseradish peroxidase (HRP) were added for 1 h at room temperature. For detection, membranes were incubated in HRP substrate (Pierce™ ECL Plus Western Blotting Substrate, Thermo Fisher Scientific) for 10 min and the chemiluminescent signal detected with the iBRIGHT imaging system (Thermo Fisher Scientific) according to the manufacturer's protocol.

**REFERENCES**:

1. Pennington HN, Lee J. 2022. Lassa virus glycoprotein complex review: insights into its unique fusion machinery. Bioscience Rep 42: BSR20211930.

2. Dambra R, Matter A, Graca K, Akhand SS, Mehta S, Bell-Cohn A, Swenson JM, Abid S, Xin D, Lewis C, Coyle L, Wang M, Bunosso K, Maugiri M, Ruiz R, Cirillo CM, Fogal B, Grimaldi C, Vigil A, Wood C, Ashour J. 2023. Nonclinical pharmacokinetics and biodistribution of VSV-GP using methods to decouple input drug disposition and viral replication. Mol Ther - Methods Clin Dev 28:190–207.
